# Wetland tree barks are dynamic hotspots for microbial trace gas cycling

**DOI:** 10.1101/2024.07.02.601631

**Authors:** Pok Man Leung, Luke C. Jeffrey, Sean K. Bay, Paula Gomez-Alvarez, Montgomery Hall, Scott G. Johnston, Johannes Dittmann, Thanavit Jirapanjawat, Tess F. Hutchinson, Nicholas V. Coleman, Xiyang Dong, Elisabeth Deschaseaux, Damien T. Maher, Chris Greening

## Abstract

Wetland tree stems have recently been shown to be a major source of methane emissions. However, the microbial communities associated within these stems (the ‘caulosphere’) and their contribution to biogeochemical cycling of methane and other compounds remain poorly understood. Here, we reveal that specialised microbial communities inhabit the bark of multiple Australian tree species and actively mediate the cycling of methane, hydrogen, and other climate-active trace gases. Based on genome-resolved metagenomics, most bark-associated bacteria are hydrogen metabolisers and facultative fermenters, adapted to dynamic redox and substrate conditions. Over three quarters of assembled genomes encoded genes for hydrogen metabolism, including novel lineages of Acidobacteriota, Verrucomicrobiota, and the candidate phylum JAJYCY01. Methanotrophs such as *Methylomonas* were abundant in certain trees and coexisted with hydrogenotrophic methanogenic *Methanobacterium*. Bark-associated microorganisms mediated aerobic oxidation of hydrogen, carbon monoxide, and methane at concentrations seen in planta, but under anoxic conditions barks could become a significant source of these gases. Field-based experiments and upscaling analysis suggested that bark communities are quantitatively significant mediators of global biogeochemical cycling, mitigating climatically-active gas emissions from stems and contributing to the net terrestrial sink of atmospheric hydrogen. These findings highlight the caulosphere as an important new research frontier for understanding microbial gas cycling and biogeochemistry.

## Introduction

Forests are among the most productive, diverse, and valuable terrestrial ecosystems on Earth ^1,2^. Their rich biota, spanning tree canopies to soil microbiota, play key roles in the global cycling of carbon and other elements. The ‘phyllosphere’ encompasses the above-ground parts of plants that microbes inhabit, including leaves, stems, and flowers. Our functional understanding of the phyllosphere has lagged behind that of the soil rhizosphere and has largely focused on leaf surfaces ^3^, while neglecting other elements such as the ‘caulosphere’ (stem and bark microbiome). The caulosphere is particularly understudied and has conventionally been considered a leached oligotrophic environment that is unfavourable for microbial life ^3^. Collectively however, the caulosphere amounts to a global surface area of ∼41 million km^2^, representing an enormous ‘bark continent’ roughly the size of North and South Americas combined ^4^. Unlike many leaves, bark is present year-round and contains diverse materials with a range of textures, thicknesses, and structures. Moreover, bark can accumulate organic molecules, water, gases, and other nutrients that may promote microbial colonization and survival ^4–6^. The bark microbiome is likely to be an important, yet vastly understudied, interface along the soil-tree-atmosphere continuum ^7–9^.

Trees are recognized for their key roles in global cycling of carbon dioxide (CO_2_) and, increasingly, methane (CH_4_). Currently, around three trillion trees remain on Earth ^10^, and the reforestation of one trillion trees is proposed in coming decades to sequester CO_2_, suggesting significant future changes in forested areas and tree biomass globally ^11,12^. Despite trees being an overall net sink of CO_2_ owing to canopy photosynthesis, tree stems were recently identified as hotspots for emission of the potent greenhouse gas CH_4_ ^13,14^. In particular, wetland tree stems can be a major source of this gas, accounting for half of all CH_4_ emissions from the Amazon floodplain, with tropical wetland trees potentially contributing ∼37 teragrams (Tg) CH_4_ yr^-1^ to the global CH_4_ budget ^15–19^. Internal sapwood CH_4_ and CO_2_ concentrations can be several orders of magnitude higher than in the atmosphere ^20–25^ as a result of plant cellular respiration and/or soil/stem microbial methanogenesis and respiration, with gases transported axially via the transpiration stream and bark pathways, then diffused radially through the caulosphere to the atmosphere (**Fig. 1A**) ^22,23,26–28^. Recent evidence based on metabarcoding suggests that microbiota associated with tree stems may play a hidden role in the production and uptake of gases. For example, methanogenic archaea (Methanobacteriales) are highly abundant in sap and heartwoods of various high CH_4_-emitting cottonwoods (*Populus sp.*) ^26,28–30^. In subtropical wetland trees, bark-dwelling methanotrophic bacteria accounted for up to 25% of the total microbial community, and actively mitigated tree stem CH_4_ fluxes to the atmosphere by ∼30 - 40% ^31,32^. These findings highlight the complex biophysical and microbial dynamics in the caulosphere that impact global atmospheric composition.

**Figure 1.**
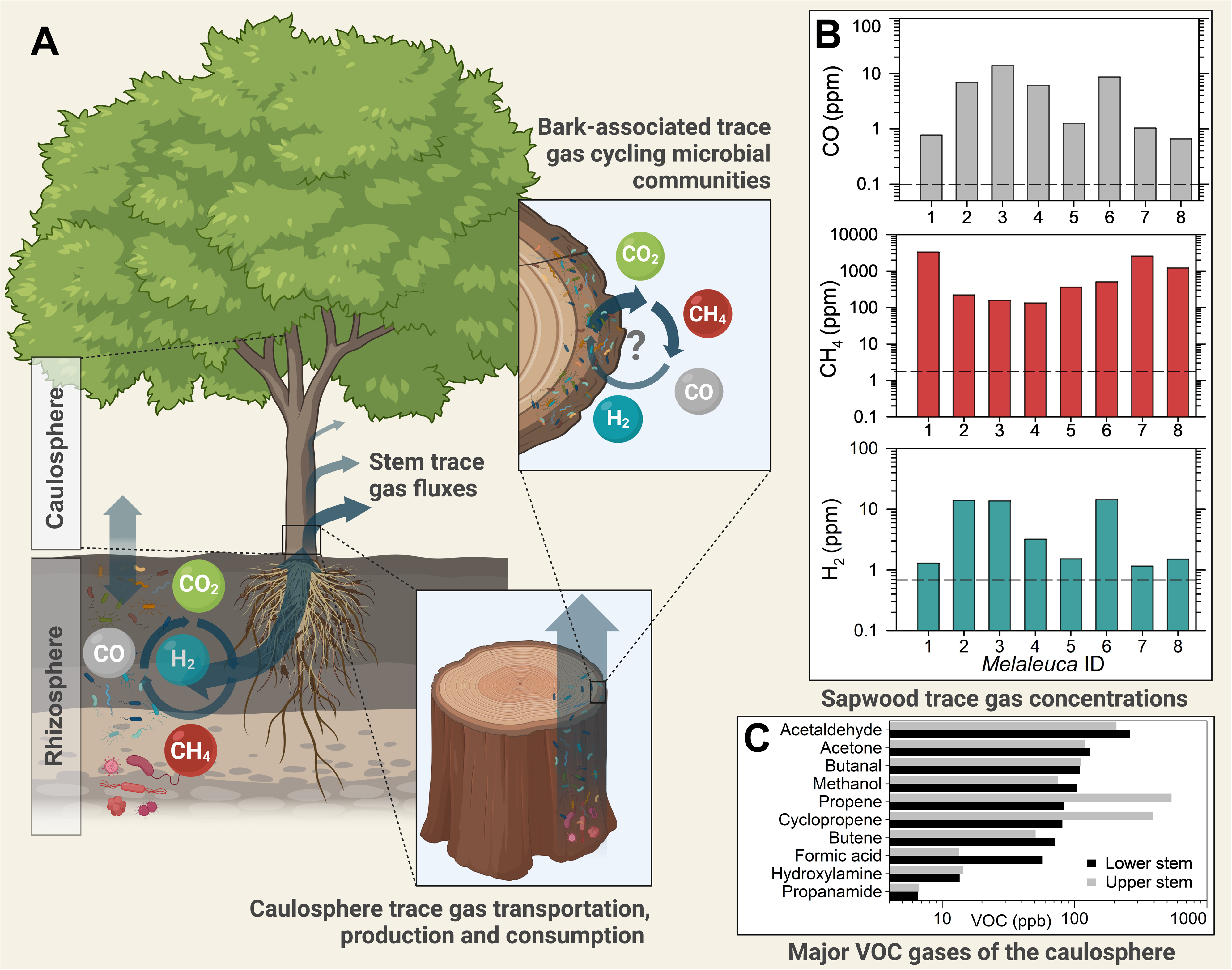
Tree stems are biogeochemical hotspots for trace gas cycling. **(A)** Conceptual diagram of the caulosphere highlighting how tree bark may be both a medium or gas transport and a niche for microbial communities adapted to trace gas cycling; **(B)** Sapwood trace gas concentrations from a subtropical wetland dominated by *Melaleuca quinquenervia* (n = 8 trees). Atmospheric levels of CO, CH_4_, and H_2_ are shown as horizontal dashed lines. Y-axis is presented in log scale; **(C)** Major VOCs emitted from tree stems (ppb) collected from a flooded *M. quinquenervia* tree at 40 and 145 cm stem heights. The X-axis is presented in log scale.

Research on the cycling of other climatically active and biologically relevant gases within forests has previously only focused on soil sources and sinks ^33^. Besides CH_4_, hydrogen (H_2_) (average 0.53 ppm) ^34^ and carbon monoxide (CO) (av. 0.1 ppm) ^35^ are the most abundant reduced atmospheric trace gases. Both H_2_ and CO are important indirect greenhouse gases, with mean global warming potentials (CO_2_-equivalent) of 12 and 2, respectively, due to their high reactivity with tropospheric hydroxyl radicals, which affect the lifetime of CH_4_ and CO_2_ ^36,37^. Biologically, these gases are fundamental energy and carbon sources for bacteria and archaea, and microbes capable of oxidizing H_2_ and CO at atmospheric concentrations are now recognized as dominant members in oxic soils, responsible for the net annual uptake of 70 Tg of H_2_ and 250 Tg of CO from the atmosphere ^33,38,39^. The activities of these trace gas oxidizers also underlie primary production in oligotrophic ecosystems such as Antarctic deserts ^40,41^ and oxic caves ^42^. In anoxic habitats, these gases are produced and used during diverse metabolic processes, including anaerobic respiration, fermentation, and methanogenesis ^43,44^. Other climatically active gases, including volatile organic compounds (VOCs) such as isoprene and short-chain alkanes, are also known to be emitted by vegetation and are metabolized by soil bacteria ^45–47^. Despite the climatic and ecological significance of trace gas cycling, it is unknown whether microbial communities within the oxic and anoxic microniches of the caulosphere may also metabolize these gases. Hence, the wider potential roles of tree stems globally, as trace gas sources and sinks, remain poorly constrained.

In this study, we integrate genome-resolved metagenomics with *in situ* and *ex situ* biogeochemical techniques to provide multiple lines of evidence for the composition, function, and activities of tree bark microbial communities. The primary field-site was dominated by ‘Broad-leaf paper bark’ trees (*Melaleuca quinquenervia*), an endemic genus representing 5% of total forest cover in Australia ^48^, and also an important invasive species introduced throughout (sub)tropical and temperate lowland regions of all continents ^49,50^. At this wetland forest, we quantified the internal concentrations of major trace gases and VOCs from tree stems, including the first measurements of H_2_ and CO. Tree bark metagenomes were collected in two consecutive years to determine the abundance, prevalence, and metabolic potentials of trace gas metabolisers. We measured metabolic activities of the caulosphere microbiota in consuming and producing CH_4_, H_2_, and CO using *ex situ* oxic and anoxic microcosm incubation experiments. *In situ* stem flux measurements and modelling across dry and wet seasons, and along axial tree stem heights, were performed to demonstrate the biogeochemical significance of these processes. Finally, the broader significance of the insights from the *Melaleuca* study was evaluated by sequencing the caulosphere metagenomes of seven other tree species, from wetland to upland forests.

## Results

### Tree stems contain elevated levels of trace gases and host numerous microbes

To quantify internal concentrations of trace gases methane (CH_4_), hydrogen (H_2_), and carbon monoxide (CO) in tree stems, we collected equilibrated gas samples from within the sapwood layer of eight paperbark trees (*M. quinquenervia*). Gas chromatography analysis confirmed that CH_4_ is present at high levels in the stems, in line with previous reports ^20–22,24,25^, and revealed for the first time that H_2_ and CO are present at elevated concentrations across all sampled trees (**Fig. 1B**). Relative to atmospheric levels, stem H_2_ (1.17 - 14.46 ppm), CO (0.66 - 13.98 ppm), and CH_4_ (136 - 3378 ppm) were on average ∼8, ∼200, and ∼600 times higher, respectively (**Table S1**). In addition, proton transfer reaction–mass spectrometry analysis (PTR-MS) showed tree stems contained a rich and diverse range of VOCs at sub-ppm levels. The most abundant stem-derived VOCs collected from the air-tight chamber set-up are (cyclo)propene (81 - 540 ppb), acetaldehyde (207 - 261 ppb), acetone (121 - 131 ppb), butanal (111 ppb), methanol (75 - 104 ppb), formic acid (13 - 15 ppb), and other undetermined alkyl compounds, for samples collected at 40 cm and 145 cm height (**Fig. 1C, Table S2**). Altogether these results suggest that tree stems are involved not only in the cycling of major greenhouse gases (CH_4_ and CO_2_), and also other climatically relevant trace gases and VOCs.

In addition to containing various trace gases, tree stems also appear to be significant habitats for microorganisms. We extracted DNA from the bark of seven *M. quinquenervia* trees, two adjacent soils, and two freshwater samples in the same *Melaleuca* wetland forest (**Fig. S1, Table S3**) and used quantitative PCR (qPCR) of 16S rRNA genes to quantify the microbial abundance. On average, bark samples contained 5.3 × 10^9^ 16S rRNA gene copies per gram of wet weight (1.8 – 8.7 × 10^9^ copies/g), comparable to surrounding soil samples (2.0 × 10^10^ copies/g) and over 200-fold higher than the adjacent surface waters (2.4 × 10^7^ copies/ml) (**Dataset S1**). The presence of an abundant community of microbes at the tree/atmosphere interface suggested they might be supported by the reduced gases present in this environment as carbon sources and/or electron donors; this hypothesis was tested next via surveying the metabolic genes present in the caulosphere metagenome.

### Unique microbial lineages enriched with gas metabolism genes reside within tree barks

Metagenomes from barks, soils, and waters were sequenced to holistically profile the composition and capabilities of their microbial communities. Community profiling using single-copy taxonomic marker genes (**Fig. 2A, Dataset S2**) and beta diversity analyses (**Fig. S2**) confirmed tree barks harbour microbial communities that are distinct from those seen in neighbouring soils and waters. Notably, *M. quinquenervia* barks were dominated by three bacterial families, Acidobacteriaceae (31.6%), Mycobacteriaceae (17.4%), and Acetobacteraceae (5.9%). We reconstructed 114 dereplicated high- and medium-quality bark metagenome-assembled genomes (MAGs) from the metagenomes, which together mapped to ∼40% of total sequences (**Fig. 2B**). Based on Genome Taxonomy Database R08-RS214 (GTDB) ^51^, all the MAGs belonged to unclassified species from ten bacterial phyla, the most abundant being genera *Terracidiphilus* (av. 23.2% in community), *Mycobacterium* (16.8%), *Acidocella* (3.1%), acidobacterial Palsa−343 (3.1%), and verrucomicrobial UBA11358 (1.8%).

**Figure 2.**
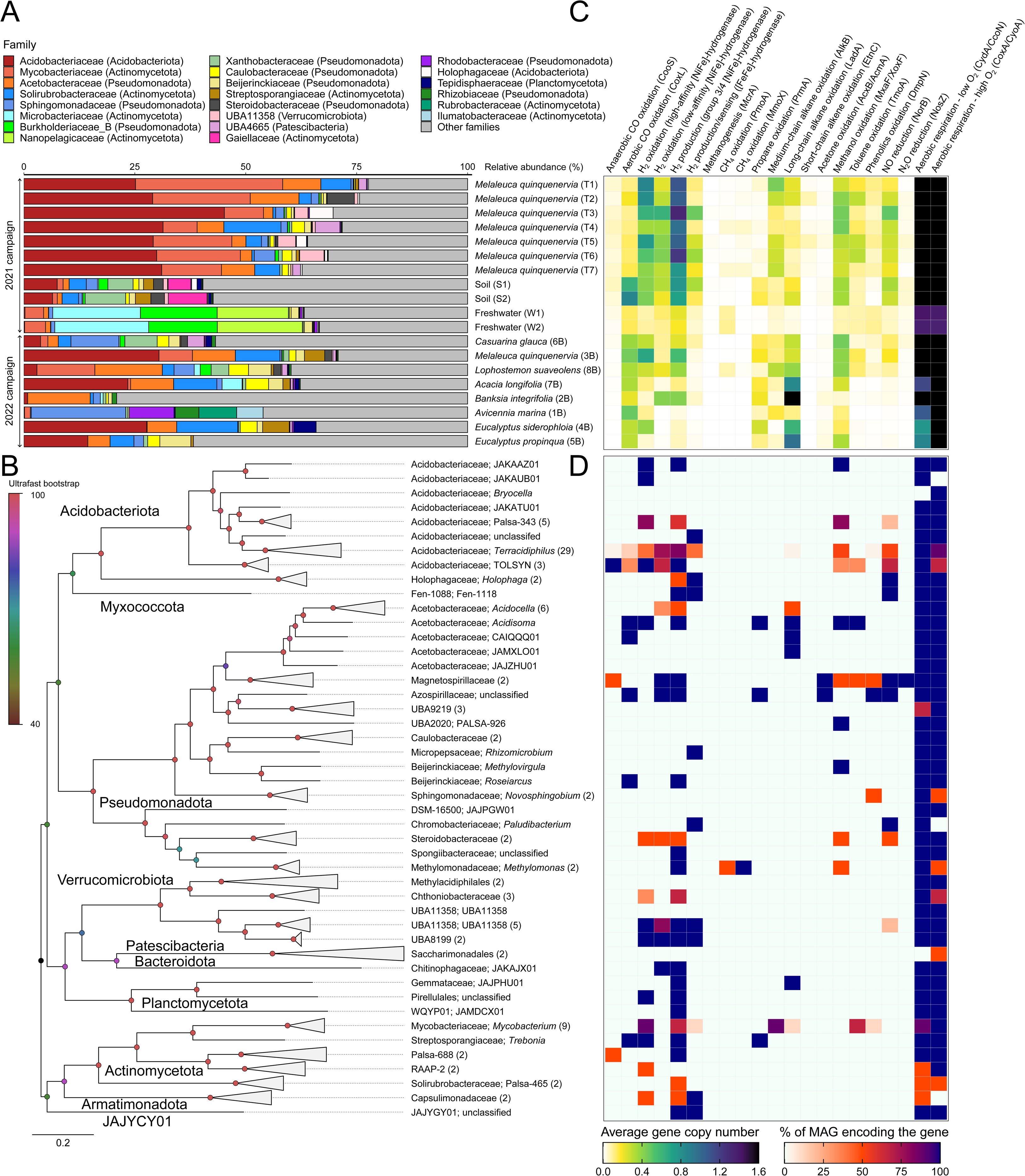
Composition and capabilities of tree bark microbial communities. (**A**) Family-level composition of microbial communities in barks of *Melaleuca quinquenervia* (n = 8 trees), surrounding wetland soils (n = 2) and freshwaters (n = 2), and barks of seven other tree species based on metagenomic reads of 59 universal single-copy marker genes. Families that do not exceed 5% abundance in any sample were grouped to “Other families”. (**B**) Maximum-likelihood phylogenomic trees of 114 medium and high-quality bacterial MAGs from *M. quinquenervia* barks based on GTDB bacterial marker genes. Leaves were collapsed at the genus level. Tip labels indicate family and genus level classification, with brackets showing the number of MAGs from each genus. Nodes are coloured based on branch support by 1000 ultrafast bootstrap replicates. For panels (**A**) and (**B**), bacterial and archaeal taxonomy is based on the GTDB release 08-RS214. (**C**) Heatmap showing average copy numbers of metabolic marker genes involved in metabolism of gases, including carbon monoxide, hydrogen, methane, VOCs (alkane, alkene, acetone, methanol, aromatics), nitric oxide, nitrous oxide, and oxygen in the total tree bark communities. (**D**) Heatmap showing the presence of the above metabolic marker genes in MAGs.

To evaluate the metabolic capacities of the bark microbes, we performed homology-based searches for marker genes involved in the cycling of gases and other compounds on metagenomic short reads (**Dataset S3**). The average copy number of the gene per organism was inferred by normalizing against reads to universal single copy ribosomal protein genes ^41^. In line with our previous report that methanotrophs are significant members of the community in this CH_4_ rich environment ^31^, particulate and soluble CH_4_ monooxygenases (*pmoA* & *mmoX*) were each present at ∼0.02 copies/organism (**Fig. 2C**). The copy number of *pmoA* in barks was on average 4.7-fold higher than in soils, whereas *mmo*-encoding methanotrophs were nearly absent in soils and waters. Methanogens coexisted with these methanotrophs within barks, soils, and waters, albeit at low abundance (av. community *mcrA* copies 0.0039, 0.0039, and 0.0027, respectively) (**Dataset S3**). Based on the sequence analysis, H_2_ was the most widely metabolized reduced gas among bark-dwelling microbes, with high-affinity uptake [NiFe]-hydrogenases mediating H_2_ oxidation at atmospheric levels (predominantly group 1h lineage) and low-affinity uptake [NiFe]-hydrogenases (represented by group 1d lineage) estimated to be present at 0.62 and 0.38 copies/organism, respectively (**Fig. 2C, Dataset S4**). This suggests microbes may efficiently harvest both elevated H_2_ from stem gas and trace H_2_ from air through distinct uptake enzymes. Remarkably, the typically O_2_-sensitive group 3 and 4 [NiFe]-hydrogenases responsible for fermentative and anaerobic respiratory H_2_ production were also encoded by large proportions of bark microbes at 0.77 and 0.30 copies per organism, respectively (**Fig. 2C**). In further testament of widespread bark H_2_ metabolism, [FeFe]-hydrogenases associated with fermentative H_2_ production and H_2_ sensing were also present at high levels (0.23 copies/organism). All the above functionally diverse hydrogenases occurred at even higher abundance in barks than in soils, which is remarkable since the latter harbour many hydrogenotrophic and hydrogenogenic bacteria and archaea ^38,44,52^.

Numerous microbes may also utilise the elevated gas substrates present in tree stem as carbon sources, electron acceptors, and electron donors. The [MoCu]-CO dehydrogenase (*coxL*) and anerobic [NiFe]-CO dehydrogenase (*cooS*) enzymes were encoded by 13.5% and 4.4% of microbes in the bark communities, respectively (**Fig. 2C**). Regarding VOC metabolism, genes encoding soluble diiron monooxygenases (SDIMOs) were the most widespread, especially lineages involved in oxidation of aromatics (*tmoA* and *dmpN*; 20.0% and 8.0%, respectively), short-chain alkenes (*etnC*; 3.0%), and short-chain alkanes (*prmA*; 2.2%). Significant proportions of the community also encoded assimilatory and oxidative pathways for methanol (*mxaF*/*xoxF*; 34.9%), medium-chain alkanes (*alkB*; 25.7%), acetone (*acxB*/*acmA*; 1.1%), as well as formic acid and ethanol (**Fig. 2C, Dataset S3**), representing VOCs detected in stem gases (**Fig. 1C, Table S2**). Interestingly, nitric oxide reduction to nitrous oxide (*norB*; 27.6%) appears to be the dominant denitrification pathway in barks, while other denitrification marker genes were encoded by less than 5% of the community (e.g. *narG*, *nirK*, *nosZ*) (**Fig. 2C, Dataset S3**).

Collectively, our metagenomic results highlight the metabolism of trace gases by bark-associated microbes as an overlooked process, with potentially substantial biogeochemical impacts. To extend and generalize this finding beyond *M. quinquenervia*, we sampled and sequenced metagenomes of seven additional tree species, including other wetland species, coastal forests, and upland forests (**Fig. S1, Table S3**). Metabolic gene profiling further confirmed a broad potential for gas metabolism in these bark communities, though the patterns appear to be strongly affected by specific environments and tree hosts. For example, bark communities from freshwater wetland trees (*Casuarina glauca* and *Lophostemon suaveolens*) were more metabolically similar to *M. quinquenervia*, characterized by their high capacity for oxidative and fermentative H_2_ metabolism, CH_4_ oxidation, and aromatic VOC degradation (**Fig. 2B**). In contrast, the bark communities of drier and well-drained upland forest trees (*Eucalyptus siderophloia* and *Eucalyptus propinqua*) were characterized by a preference for long-chain alkane degradation, but reduced capacity for anaerobic trace gas oxidation and denitrification, whereas the metabolic potentials of communities from littoral and coastal forests were intermediate between the wetland and upland sites (**Fig. 2B**). Future studies should explore environmental and biological factors that control the composition and function of bark communities.

### The most abundant bark microbes are facultative fermenters that oxidize, produce, and sense H_2_

In-depth metabolic profiling of MAGs and unbinned assemblies was performed to identify key microbial determinants and potential roles of the widespread trace gas metabolism. Bark communities were broadly adapted to the varying O_2_ availability and unique substrate conditions found in tree stems. Most MAGs co-encode canonical terminal oxidases (cytochrome *aa_3_*and *bo_3_* types; *coxA*/*cyoA*) for aerobic growth and high-affinity terminal oxidases (cytochrome *bd* and *cbb_3_*types; *cydA*/*ccoN*) to support O_2_ scavenging under microoxic conditions (**Fig. 2D**). Under O_2_ limitation, microbial growth is likely driven by fermentation, as tree stems are carbon-rich yet oxidant-restricted environments ^14,53^. This is reflected by the overrepresentation of genes involved in fermentation (especially formatogenic, acetogenic, solventogenic, and lactogenic pathways), while the capacity to respire inorganic substrates such as via denitrification, sulfate reduction, iron reduction, and reductive dehalogenation is scarce (**Fig. S3, Dataset S4**).

In addition to being abundant overall, the H_2_-metabolizing microbes in barks were also taxonomically diverse, with 89 out of 114 MAGs from 27 families encoding at least one copy of [NiFe]- or [FeFe]-hydrogenase (**Fig. 3, Dataset S4**). Phylogenetic analysis revealed that dominant bacterial species represented by high-quality genomes often encode multiple hydrogenases of distinct subgroups for high-affinity uptake (groups 1h, 1l, 1f, 2a), low-affinity uptake (groups 1a-e), fermentative production (groups 3b, 3d), bifurcation (group A3), and sensing (groups C3, 2b-c) of H_2_ (**Fig. 3**) ^54,55^. These span the most abundant genera, including acidobacterial lineages *Terracidiphilus* (1d, 1h, 3d, A3, C3), Palsa-343 (1f, 3d), and TOLSYN (1c, 1h, 3d, 4c, 4g), verrucomicrobial lineages UBA11358 (1d, 1h, 3d, A3, C3) and UBA8199 (1d, 1h, 3d, 4e, A3, C3), actinobacterial lineage *Mycobacterium* (1h, 2a, 3b/3d), and the novel phylum JAJYGY01 (3d, 4e, A3, C3). Through this arsenal of hydrogenases, these microbes are likely to be able to flexibly adapt their metabolism to maintain energy needs and redox balance in response to the dynamic redox conditions and H_2_ concentrations within tree stems. For example, in the case of the ubiquitous genus *Terracidiphilus*, the co-occurrence of group 1d and 1h [NiFe]-hydrogenases would enable them to efficiently oxidize H_2_ present at high concentrations in stem gas and at low concentrations from the atmosphere, inputting electrons to terminal oxidases for energy conservation under oxic conditions (**Fig. 2D**). Under anoxic conditions, the same bacterium could switch to fermentation, using the group 3d [NiFe]-hydrogenase to dispose of excessive reductants as H_2_; this is consistent with our pure cultures studies that show mycobacteria adapt to hypoxia by switching from aerobic respiration to fermentation ^56,57^. The sensory group C3 [FeFe]-hydrogenase likely regulates hydrogenase expression in response to H_2_ accumulation under anoxic conditions, whereas the H_2_ bifurcating/confurcating group A3 [FeFe]-hydrogenase likely acts as a redox valve, recycling cofactors under anoxic conditions. These microbes and hydrogenases similarly predominate in the bark metagenomes of *C. glauca* and *L. suaveolens*, alluding to a common adaptation within wetland-associated tree species (**Fig. 2C & S4**).

**Figure 3.**
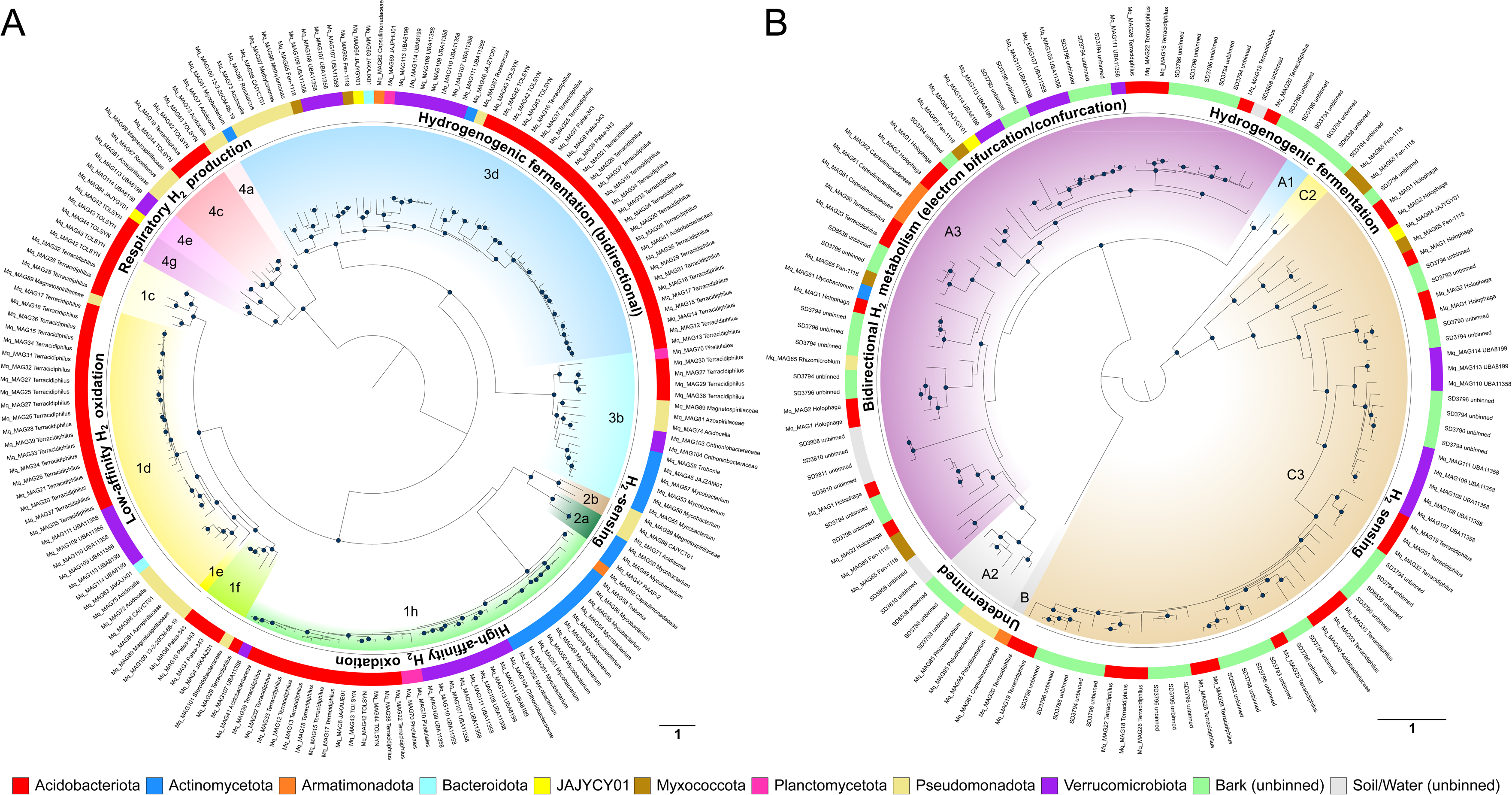
Numerous bark-associated microbes use diverse hydrogenases to shift between respiration and fermentation. Maximum-likelihood phylogenetic reconstructions of the amino acid sequences of (**A**) [NiFe]-hydrogenases from MAGs and (**B**) [FeFe]-hydrogenases from MAGs and unbinned contigs dereplicated at 97% identity. Trees were rooted at mid-point, nodes with >75% branch support (1000 ultrafast bootstrap replicates) are shown in black dots, and the scale bar indicates the average number of substitutions per site. Hydrogenase subgroups and functions were classified based on the hydrogenase database (HydDB) ^55^. The coloured outer ring denotes phylum-level taxonomy of MAGs and environmental origins of unbinned hydrogenase sequences.

While H_2_ metabolism is a generalist trait of the caulosphere, the oxidation of the most abundant reduced gas, CH_4_, is a specialist one. Two methanotroph MAGs belonging to unclassified *Methylomonas* were recovered, with a near complete MAG (∼100%; Mq_MAG97) from this genus encodes a single copy each of the *pmo*, *mmo*, and the *pmo*-like *pxm* operon (**Dataset S4**). This MAG also encodes a group 3d [NiFe]-hydrogenase, which may support their survival under anoxia. Several unbinned *pmoA* and *mmoX* sequences from *M. quinquenervia* bark also cluster within the *Methylomonas* clade (**Fig. 4A-B**), in agreement with their previously reported high abundance ^31^. Other *Methylomonas pmoA* sequences were identified in *L. suaveolens* bark and freshwater, although the methanotrophic populations in these locations were distinct, with *Methylocystis* being dominant in the former and diverse methanotrophs present in the later (**Fig. 4A**). Interestingly, *mmo*-encoding methanotrophs were scarce outside *M. quinquenervia* bark (**Fig. 2C**). Oxidative *mcrA* (*r-mcrA*) was not detected in any sample, suggesting a low capacity for anaerobic methane oxidation in bark communities (**Dataset S3**). We recovered two high-quality MAGs representing two mesophilic families of Methylacidiphilales, yet neither encodes CH_4_ monooxygenase nor other key methanotrophy genes (**Fig. 2D**). Thus, contrary to our initial prediction and unlike other members of this order ^31^, it is unlikely that these bacteria contribute to CH_4_ oxidation in bark.

**Figure 4.**
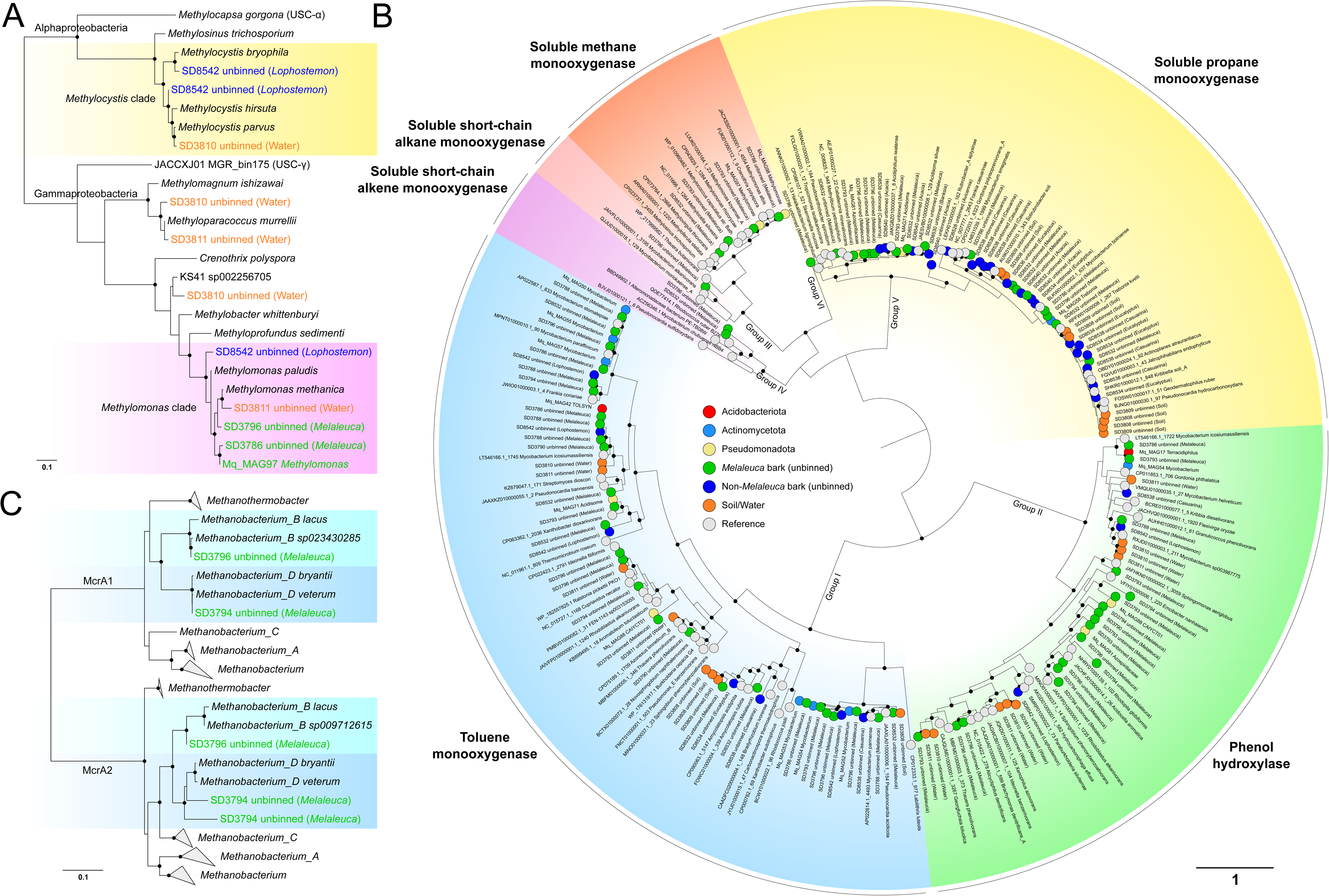
Diversity and distribution of enzymes for methane and VOC metabolism in bark microbial communities. (**A**) Particulate CH_4_ monooxygenase (PmoA; aerobic CH_4_ oxidation marker), (**B**) soluble diiron monooxygenase (TmoA, DmpN, MmoX, EtnC, PrmA; aerobic VOC metabolism markers), and (**C**) methyl-CoM reductase (McrA; methanogenesis marker) Maximum-likelihood phylogenetic trees were constructed from reference sequences, MAGs, and unbinned contigs dereplicated at 97% identity. Trees were rooted at mid-point, nodes with >75% branch support (1000 ultrafast bootstrap replicates) are shown in black dots, and the scale bar indicates the average number of substitutions per site. SDIMO subgroups and functions were classified based on previous studies ^58,59^, but it is important to note that the common names shown (e.g. ‘phenol hydroxylase’) are indicative rather than rigorous descriptions of their substrate preferences. Coloured circles at the leaf nodes in panel (**B**) denote phylum-level taxonomy of MAGs and environmental origins of unbinned monooxygenase sequences.

Diverse taxa are predicted to mediate the metabolism of other trace gases and VOCs within tree barks. CO oxidation genes were found among Acidobacteriota, Actinobacteriota, and Proteobacteria (**Fig. 2D**). Phylogenetic analysis of CoxL revealed actinobacterial and proteobacterial clades predominate aerobic CO oxidation, with the mixed 1 clade represented by *Terracidiphilus* sequences also abundant (**Fig. S5**). Six MAGs (*Terracidiphilus*, TOLSYN, JAJZAM01, Magnetospirillaceae) encode *cooS* (**Fig. 2D**), five of which encode a group 4 [NiFe]-hydrogenase / CO dehydrogenase supercomplex, known to couple anaerobic CO oxidation to respiratory proton reduction ^60,61^ (**Fig. 3A**). Thus, H_2_ and CO metabolism are likely to be coupled in tree bark. *Mycobacterium* was inferred to be the most dominant genus in oxidation of medium-chain hydrocarbons and aromatics (**Fig. 2D**), consistent with previous literature ^62,63^. Proteobacterial MAGs from *Acidisoma*, Magnetospirillaceae, and Azospirillaceae were also enriched with a repertoire of genes for oxidation of acetone, short-chain alkanes, and toluene/phenol derivatives (**Fig. 2D**). Additionally, a wide diversity of monooxygenases for linear and aromatic hydrocarbon oxidation were identified in unbinned sequences (**Fig. 4B**). Methanol metabolism is widespread among Acidobacteriota and Proteobacteria. Notably, calcium-dependent methanol dehydrogenase (*mxaF*) is only encoded by *Methylomonas* and *Methylovirgula* MAGs, while other MAGs encode the lanthanide-dependent enzyme (*xoxF*) (**Dataset S4**). Finally, most of the bark-associated bacteria were predicted to consume and produce other VOCs such as formate, acetate, acetaldehyde, and ethanol as part of their central carbon metabolism (**Fig. S3**).

### Bark microbes consume and produce trace gases rapidly in response to dynamic redox conditions

To confirm these genomically-derived hypotheses, we prepared microcosm assays on both *M. quinquenervia* and *C. glauca* bark samples to test whether the associated microbes metabolized the major trace gases aerobically and/or anaerobically. In the oxic experiment, barks were incubated in closed vials with ambient air headspace supplemented with 10 ppm each of H_2_, CO, and CH_4_ (H_2_ and CO resembling their concentrations within stem gas). The consumption of trace gases was monitored by ultra-sensitive gas chromatography over several days (**Fig. 5A-C**). In line with the high abundance of aerobic H_2_ oxidizers inferred from metagenomics (**Fig. 2**), H_2_ was rapidly consumed by all tested barks within 24 hr (-46 ± 3 to -74 ± 13 nmol/g of bark/d in *M. quinquenervia* and *C. glauca* bark, respectively) (**Fig. 5A, Table S4**). CO was also consumed by bark communities, but much more slowly than H_2_ oxidation (8 – 23-fold slower; **Table S4**). The *C. glauca* barks harboured three-fold more CO oxidizers than *M. quinquenervia* barks, and also showed faster CO consumption (**Fig. 5B**). These experiments uncover that barks are active sinks for the climate active trace gases H_2_ and CO.

**Figure 5.**
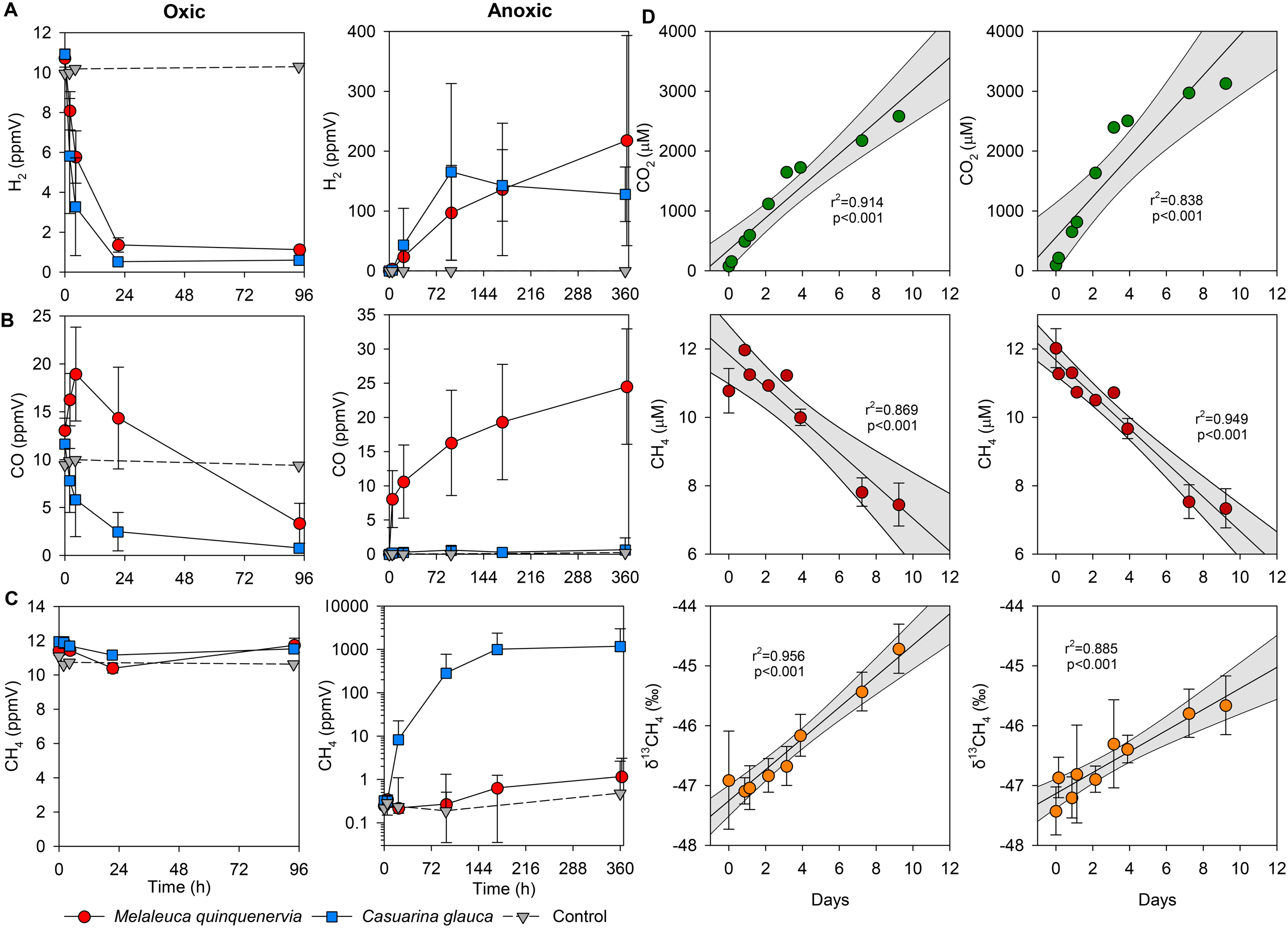
Metabolism of hydrogen, carbon monoxide, and methane in oxic and anoxic microcosms. (**A-C**) Microcosms of *Melaleuca quinquenervia* barks (red circle) and *Casuarina glauca* barks (blue squares) (n = 3 trees) were incubated at 20°C, with empty vials as negative controls (grey triangle with dashed line). Oxic incubations were set up with approximately 10 ppm each of H_2_, CO, and CH_4_ in the ambient air headspace while anoxic incubations were established by purging headspace with ultra-pure N_2_. Data are presented as mean ± S.D. values of headspace (**A**) H_2_, (**B**) CO, (**C**) CH_4_ mixing ratios. (**D**) Cavity ring-down spectroscopy measurements of CH_4_ oxidation to CO_2_ in *M. quinquenervia* barks (n = 2 trees). These were oxic microcosms containing ambient air spiked with CH_4_ at ∼500 ppm, incubated at 21°C. Plots show CO_2_ production (green dots), CH_4_ consumption (red dots) and δ^13^C-CH_4_ (‰) enrichment (orange dots) over time. Data are presented as mean ± S.D. values. Grey areas are 95 % CI.

No CH_4_ consumption was detected when barks were incubated at near atmospheric levels (∼10 ppm) (**Fig. 5C**), indicating a minimal capacity for high-affinity CH_4_ uptake. We therefore carried out a separate oxic incubation using a high CH_4_ concentration (∼500 ppm) similar to stem *in situ* concentrations (**Fig. 1B**) and observed CH_4_ consumption, CO_2_ production, and changes in the carbon stable isotope ratio (δ^13^C) of CH_4_, linked to fractionation by methanotrophic activity. At these concentrations, the bark showed CH_4_ oxidation at rates of -4.5 to -5.6 nmol/g of bark/d (**Fig. 5D, Table S5**), suggesting CH_4_ uptake activity was mediated by low-affinity methanotrophs. Evidence that the observed CH_4_ oxidation was due to biological processes was seen in the corresponding increase in δ^13^C of the headspace CH_4_ (**Fig. 5D**). These observations echo with the dominance of known low-affinity methanotrophs in the community (e.g. *Methylomonas, Methylocella;* **Fig. 4A-B**) and our earlier report that bark-mediated CH_4_ oxidation plateaued at 20 – 80 ppm ^31^. Taken together, the results indicate that wetland barks are unlikely to directly oxidize atmospheric CH_4_, but methanotrophs do help moderate the net emissions of CH_4_ from the caulosphere in wetland environments.

In contrast to oxic conditions, barks can become a source of these trace gases under anoxic conditions. In anoxic microcosms, H_2_ was produced by barks from both *M. quinquenervia* and *C. glauca* at substantial rates (43 ± 20 to 70 ± 35 nmol/g of bark/d, respectively) (**Fig. 5A, Table S4**), mostly likely reflecting high microbial fermentation activities. CO was slowly produced by *M. quinquenervia* barks (7 ± 2 nmol/g of bark/d), whereas there was negligible CO production from *C. glauca* barks (**Fig. 5B, Table S4**). CO emissions were probably due to abiotic decomposition of carbon-rich materials, as previously demonstrated in plant litter and soils ^64–66^. Further testing is needed to determine if the differences in CO emissions between the two species of tree were due to variations in the chemical recalcitrance of carbon compounds to abiotic breakdown or microbial activities such as CO-dependent hydrogenogenesis, acetogenesis, or methanogenesis. Unexpectedly, despite the low abundance of methanogens, methanogenesis occurred at substantial rates in *C. glauca* barks, with up to 1000 ppm CH_4_ produced after four days of incubation (av. CH_4_ production of 120 ± 117 nmol/g of bark/d) (**Fig. 5C**). CH_4_ production also occurred in *M. quinquenervia* bark, but at a rate several orders of magnitude lower (0.01 ± 0.1 nmol/g of bark/d). While no archaeal MAG was recovered, sequences of *mcrA* related to *Methanobacterium lacus* and *Methanobacterium bryantii* were assembled from *M. quinquenervia* bark metagenome (**Fig. 4C**), along with group 3a, 3c, 4h, and 4i [NiFe]-hydrogenases associated with this same genus (**Fig. S4, Dataset S5**). *Methanobacterium* species are known from previous work to produce CH_4_ using H_2_ and CO ^67^ and thus they are likely to be the dominant methanogens in *M. quinquenervia* and *C. glauca* (this conclusion is also supported by PhyloFlash-based community profiling; **Dataset S6**). Hydrogenotrophic methanogenic activity may explain the increase in CH_4_ but the early leveling off of H_2_ production and the minimal CO accumulation in anoxic *C. glauca* bark (**Fig. 5A-C**).

A concordant pattern of activity to that described above was observed for barks collected at another sampling time (**Fig. S6**) and during a second microcosm assay containing *M. quinquenervia* bark collected from 0.4, 2.0, 6.0 and 8.8 m stem heights (**Fig. S7, Table S3 & S6**). The latter experiment revealed similar uptake rates between lower and upper bark for H_2_ (-74.1 to -87.4 nmol/g of bark/d) and CO (-4.1 to -7.7 nmol/g of bark/d) under oxic conditions; while under anoxic conditions, the production of H_2_ (2.21 to 0.45 nmol/g of bark/d) and CO (3.84 to 1.95 nmol/g of bark/d) decreased with stem height. A large spike of CH_4_ production was observed in the bark collected from 2.0 m height (193 nmol/g of bark/d) (**Table S6**). These results further validate that the observed activities were not an isolated phenomenon. Overall, the data show that diverse trace gas metabolizing microbes co-exist in tree stems throughout the height of the tree, and their rapid activity is likely to contribute to the regulation of trace gas fluxes from forested ecosystems.

### Tree stems are biogeochemical hotspots of trace gas cycling

To further explore the wider biogeochemical and ecosystem implications of *in situ* trace gas cycling within tree stems, we monitored and compared gas fluxes from eight *M. quinquenervia* trees during dry and wet seasons (**Fig. 6 & S8**). Gas samples collected from static chambers installed on stems were analysed for CH_4_, CO_2_, H_2_, and CO through a combination of cavity ring-down spectrometry and gas chromatography. The wetland tree stems showed net emissions for CO_2_, CH_4_, and CO, but were consistently a net sink for H_2_. The overall pattern was consistent for all trees across both seasons, though the magnitude of fluxes for CH_4_, CO and H_2_ was significantly affected by seasonality (*p* < 0.05, Welch’s *t* test) **(Fig. 6A, Table S7**). Trees emitted an average of 80.5 ± 18.5 µmol m^-2^ d^-1^ of CO_2_ during the dry season, which increased to 123.8 ± 23.1 µmol m^-2^ d^-1^ during the wet season (**Fig. 6, Table S7**). Dry season tree stem CH_4_ emissions were 155.3 ± 72.7 µmol m^-2^ d^-1^and significantly increased (*p* < 0.001, Welch’s *t* test) by 65-fold during the wet season (**Fig. 6B, Table S7**). While prior studies have revealed that the majority of CO_2_ and CH_4_ emissions are derived from subsoil and/or host respiration ^22,23,68,69^, activities of microbial communities in stems likely also enhanced their production and emissions. Interestingly, whereas bark communities can consume CO, there was a net positive flux of this gas at the stem surface at a comparable magnitude to CO_2_ (7.37 ± 1.11 µmol m^-2^ d^-1^ in dry conditions and 16.1 ± 2.75 µmol m^-2^ d^-1^ in wet conditions) (**Fig. 6, Table S7**). This suggests abiotic CO production and/or that the strong soil source of CO is being transported upwards via stem and bark layers at rates exceeding bark-associated microbial CO oxidation.

**Figure 6.**
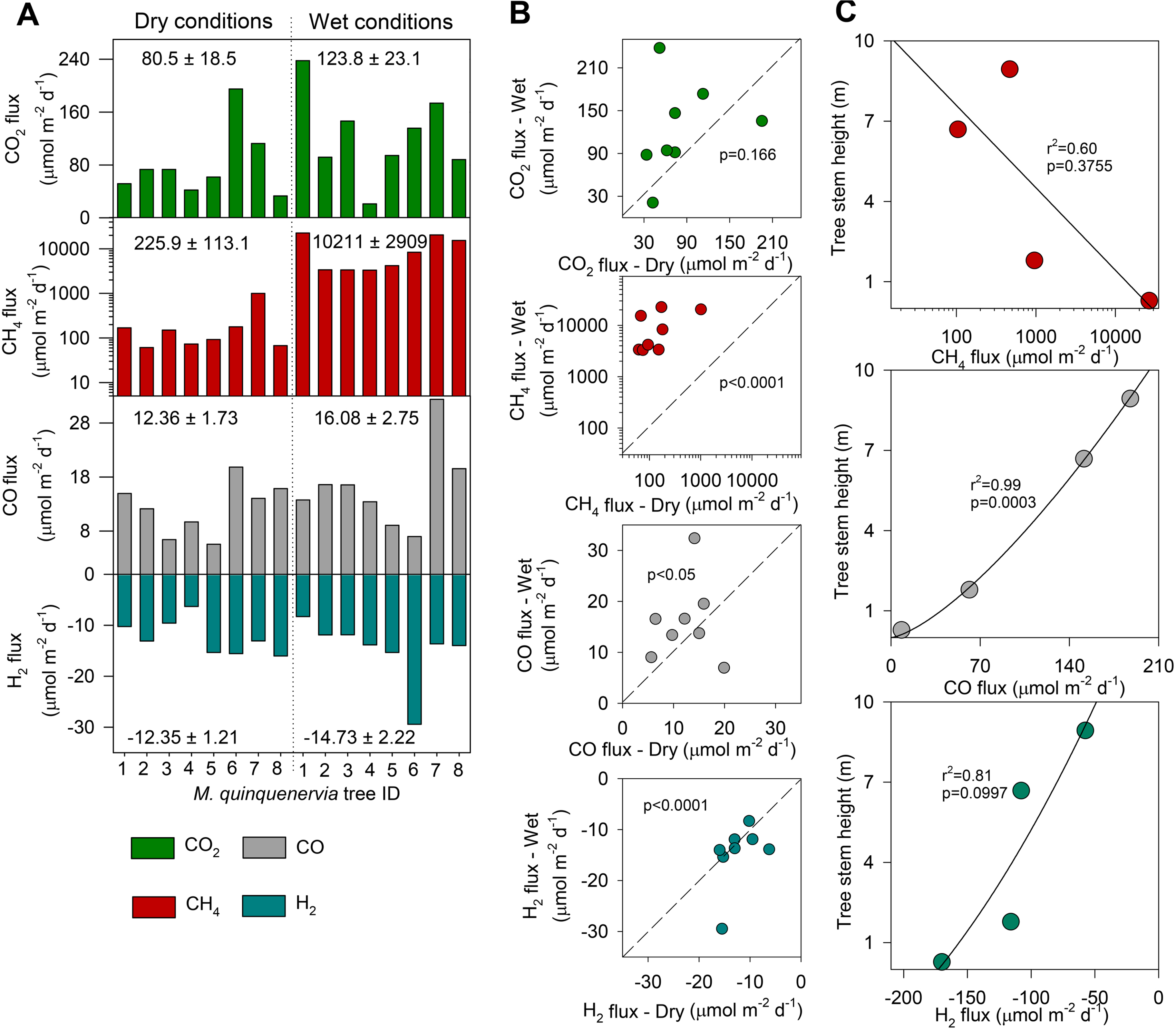
Tree stems mediate substantial trace gas fluxes. **(A)** Tree stem CO_2_, CH_4_, CO and H_2_ trace gas fluxes measured from the lower stems of wetland *Melaleuca quinquenervia* (n = 8 trees) during dry (**left**) and wet (**right**) seasonal conditions; **(B)** Correlations of paired wet vs dry stem trace gas fluxes showing significant differences for CH_4_, CO, and H_2_ (*p* < 0.05, Welch’s *t* test) between fluxes of each field campaign. The dashed line shows a 1:1 mixing ratio, i.e., same values in both campaigns; **(C)** Regressions of trace gas fluxes vs axial tree stem height in *M. quinquenervia*. Note: Non-significant difference for CO_2_ fluxes (*p* = 0.166) between campaigns in Fig. 6B; all rates are in µmol m^-2^ (of stem) d^-1^.

Finally, tree stems were associated with a net *in situ* uptake of atmospheric H_2_, with activities in the wet season (-15.5 ± 2.22 µmol m^-2^ d^-1^) significantly higher than in the dry season (-5.79 ± 0.82 µmol m^-2^ d^-1^) (**Fig. 6, Table S7**). Furthermore, the *in situ* H_2_ consumption was also observed in barks at all sampling heights up to near the tree canopy (9 m above ground) (**Fig. 6C & S7, Table S6**). In conjunction with the paired microcosm experiments (**Fig. 5**), these findings provide further evidence that bark-dwelling communities are metabolically active throughout the caulosphere. The areal rates of tree stem H_2_ consumption upscaled to the forest scale (-10.9 to -22.9 µmol m^-2^ of forest d^-1^) are lower than previously reported soil H_2_ consumption (-1853 to -2281 µmol m^-2^ of soil d^-1^ in wetland and forest soils, respectively) ^38,52^, however our results still suggest that the caulosphere may represent an overlooked and quantitatively significant biological sink of this gas.

## Discussion

Through an integrated experimental approach with systematic profiling of microbial abundance, communities, metabolic functions, and biogeochemical activities, our study has revealed the caulosphere as an overlooked microbiological and biogeochemical hotspot for important trace gases. Bark surfaces serve as both interfaces and conduits where trace gases are actively cycled between the atmosphere and the tree. Selection for trace gas metabolism in turn enables abundant and unique microbial communities to thrive within the caulosphere. Under oxic conditions, diverse caulosphere bacteria consume H_2_, CO, and VOCs from both stem gas and the atmosphere for energy conservation through aerobic respiration; while under O_2_ constraint, the community can switch to production of H_2_ and various VOCs (e.g., formate, acetate, ethanol) to sustain fermentative growth. The most ubiquitous microbes in bark appear to be those with flexible gas metabolism, enabling them to rapidly respond to changing redox conditions and substrate availability. Members of the highly abundant *Terracidiphilus* genus are remarkable examples of this versatility, and can oxidize, fermentatively evolve, bifurcate, and sense H_2_, in addition to NO respiration, CO consumption, and aerobic oxidation and fermentative production of VOCs. The unique bark environment also allows the co-existence of metabolic specialists such as low-affinity methanotrophs (e.g., *Methylomonas*) that feed on stem-enriched CH_4_ and methanogens (e.g., *Methanobacterium*) that produce CH_4_ from H_2_ and CO.

The activity of caulosphere microbial communities clearly modulates trace gas fluxes at tree stems, with dual biogeochemical roles as both consumers and producers of various climatically-active gases, including CH_4_, H_2_, CO, CO_2_, and VOCs. Laboratory measurements showed that the bark communities consume CH_4_, H_2_, and CO aerobically at significant rates that are comparable to soils. In the field, tree stems appear to be net sinks of atmospheric H_2_ across both wet and dry seasons, despite supra-atmospheric levels of this gas within sapwood. This indicates remarkable activity of high-affinity hydrogenotrophs, in line with their high cell counts in tree barks. In contrast, wetland trees emerge as a net source for CO and CH_4_, concomitant with the high concentrations of these gases within stems, but it must also be noted that numerous aerobic bark microbes are likely to be mitigating a large fraction of these emissions. Depending on environmental conditions and future climate, the metabolic flexibility of microbial communities in caulosphere indicates a potential to become a significant source of these three gases. For example, if conditions change from oxic to anoxic, methanogenesis and H_2_ production due to fermentation may be rapidly activated, whereas net CO uptake decreases. Hypoxia can be driven by increased precipitation events, and elevated host and microbial respiration; evidence for the former was seen in the significant increases in CH_4_ and CO stem fluxes from paperbark trees during the wet season. Together, these results extend recent observations that tropical wetland trees are globally significant CH_4_ hotspots, by showing that tree stems can also consume and emit other important climate-active gases. This emphasises that trees they are not just ‘passive pipes’ for gaseous transport but also host active and diverse microbes that metabolize various trace gases.

This study does not exhaustively quantify biogeochemical fluxes and microbial communities across the diversity of forest ecosystems, and indeed, our data show the high variability of trace gas fluxes across tree individuals and species, and also the strong impact of environmental conditions (wetland vs. drier upland) on these phenomena. The current study does, however, provide definitive evidence that tree stems, often covering extensive areal surface in wetland forests, are overlooked hotspots for diverse trace gas metabolism; in addition to providing the first-order activity estimates of these processes and the first data on the identities and genomes of the microbes involved. Tree stems currently represent an unaccounted for sink and source in various global greenhouse gas budgets and models, despite processing and mediating considerable biogeochemical fluxes. Ongoing efforts will be required to determine the drivers of the distribution, lifestyle, and magnitude of the activities of microbial communities in different tree hosts and environments, in addition to identifying the relative contributions of abiotic and biotic factors that control trace gas fluxes at tree stems, and importantly how climate change is influencing these processes. Resolving these knowledge gaps will have important ramifications for understanding the symbiotic roles of caulosphere microbial communities, and informing optimal forest management strategies that reinforce the sinks and minimize the sources of undesirable greenhouse gases.

## Methods

### Field sampling and bark collection

The field sites were located within subtropical north east New South Wales, Australia. The first bark samples were collected 20/5/2020 from flooded *Melaleuca quinquenervia* forest. The qPCR results targeting methanotrophs were previously published in Jeffrey *et al*. 2021 ^31^ and the metagenomes were later sequenced for this study (see methods below). A second bark collection for genomic sequencing focused on eight widely distributed species from contrasting forest biomes, during wet but non-flooded conditions; this was conducted on 10/5/2021. This included bark from trees situated in upland forests (*Eucalyptus propinqua*, *Eucalyptus siderophloia*), coastal dunes (*Banksia intergrifolia, Acacia longifolia*), wetland forests (*Casuarina glauca, Lophostemon suaveolens, M. quinquenervia*), and mangrove ecosystems (*Avicennia marina*). On 21/2/2022, tree stem bark VOC composition was determined on one flooded *M. quinquenervia* from two stem heights (methods below). For a trial microcosm experiment, a bark swatch was collected from one *M. quinquenervia* and one *C. glauca* on 3/11/2021; this was repeated on 21/1/2022 for the final microcosm experiment (methods below). *In situ* stem fluxes of H_2_, CO, CO_2_, and CH_4_, as well as sapwood stem gas composition, were measured on eight *M. quinquenervia* individuals on 14/11/2023 (dry - Spring) and the same trees were re-sampled on 6/2/2024 (flooded – Summer) to determine differences in trace gas concentrations and fluxes between hydrological seasons (methods below). *In situ* trace gas stem fluxes were later measured from multiple stem heights of 0.4 m, 2.0 m, 6.6 m and 8.8 m above the soil, on one *M. quinquenervia* on 22/3/2024. Bark swatches were also collected from the same axial heights for microcosm experiments (see below). Bark samples from two *M. quinquenervia* were collected on 20/5/2024 to conduct low-affinity CH_4_ oxidation microcosm assays.

All barks were collected using sterile methods, including a blade, gloves, and 70% ethanol to clean equipment between each sample. The bark swatches (10 × 10 cm) were stored in pre-sterilised foil pouches and kept cool until further analysis, as described in Jeffrey *et al*., 2021 ^31^. If a portion of each bark was required for DNA sequencing, it was immediately frozen upon returning to the lab (-80°C). The remaining portion of fresh barks were used for microcosm assays within a couple of days of collection. Bark volumetric moisture content (VWC %) was determined by wet weighing, then oven drying samples at 80°C to constant weight. All ancillary details are summarised in **Table S3**.

### DNA extraction and quantitative PCR

Approximately 0.1 – 0.2 g of frozen bark samples were homogenized in liquid nitrogen using a sterile pestle and mortar. DNA was extracted from the ground bark samples using the Synergy 2.0 Plant DNA extraction kit (OPS Diagnostics), following manufacturer’s instructions. Two blank extractions were prepared as negative controls. DNA from soil (∼0.25 g wet weight) and freshwater samples (50 ml filtered onto sterile 0.2µm filter papers) was extracted using the DNeasy PowerSoil Kit (Qiagen), according to manufacturer’s instructions. The purity and yield of DNA extracts were assayed by a NanoDrop ND-1000 spectrophotometer (Nanodrop Technologies Inc) and a Qubit Fluorometer (ThermoFisher Scientific), respectively. Total microbial abundance was estimated by quantifying the total 16S rRNA gene copy number through quantitative PCR targeting the V4 region using 515F (5’-GTGYCAGCMGCCGCGGTAA-3’) and 806R (5’-GGACTACNVGGGTWTCTAAT-3’) primers. Each 20 µl PCR reaction contained 1 µl of 1/10 diluted template DNA, 10 µl of 2 x LightCycler 480 SYBR Green I Master (Millennium Science), and a pair of primers at 0.2 µM each. The PCR was carried out in the Applied Biosystems QuantStudio 7 (ThermoFisher Scientific) using by the following program; 3 min at 95°C, and then 50 cycles of 30 s at 95°C, 30 s at 54°C, and 24 s at 72°C. Plasmid standards containing a copy of the *Escherichia coli* 16S rRNA gene were synthesized by GeneArt (ThermoFisher Scientific). Prior to qPCR, the plasmid standard was linearised using SpeI restriction enzyme (New England Biolabs) and serially diluted. Non-specific binding and amplification were tracked using the melt curve which initiated at the end of the PCR stage with the following program; 15 s at 95°C, 1 min at 60°C, and 15 s at 95°C for data collection.

### Metagenomic sequencing and quality control

For metagenomic sequencing, DNA was shipped on dry ice to the Australian Centre for Ecogenomics (ACE), University of Queensland. Metagenomic shotgun libraries were prepared using the Nextera XT DNA Library Preparation Kit (Illumina) and passed to paired-end sequencing (2 × 150 bp) on a NovaSeq6000 platform (Illumina). On average, sequencing yielded 37,450,000 read pairs for each of the 19 metagenomes (**Table S1**). Minimal contamination from DNA extraction and sequencing processes were evidenced by three-order magnitudes lower read pairs and 16S rRNA gene copies in the two negative controls compared to the samples. Raw metagenomic reads were quality processed through the BBDuk function of the BBTools v39.01 (https://sourceforge.net/projects/bbmap/). The rightmost base known to contain a high error rate, sequencing adapters (k-mer size of 23 and hamming distance of 1), PhiX spike-in (k-mer size of 31 and hamming distance of 1), bases from 3’ ends with a Phred score below 20, reads matched to human genomes using removehuman.sh, and resultant reads with lengths shorter than 50 were sequentially removed. Read quality before and after processing was assessed using FastQC v0.11.7 (http://www.bioinformatics.babraham.ac.uk/projects/fastqc/). 83.5% high-quality read pairs were retained for downstream analysis.

### Community analysis

Microbial community structures were determined independently using both universal single copy ribosomal protein genes and small subunit rRNA (SSU rRNA) gene in metagenomes. SingleM v.1.0.0beta7 ^70^ was used to map quality-filtered reads to fixed windows of conserved single copy ribosomal marker genes and the matched sequences were clustered to operational taxonomic units (OTUs) at 97% identity. The latest GTDB package at the time of analysis (R08-RS214; https://zenodo.org/record/5523588#.Yl6OANpByUk) was used to assign taxonomy to mapped reads. Use of the same GTDB database release for both community and MAG analysis enabled taxonomic consistency. Reads assigned to eukaryote and root only were filtered. PhyloFlash v.3.4.2 ^71^ was used to screen quality-filtered reads for the SSU rRNA sequences and assembled with the option –almosteverything. The SSU Ref NR99 database from the SILVA release 138 ^72^ served as the reference for the sequence searching and taxonomy assignment of SSU reads to the Nearest Taxonomic Units (NTUs). Full lists of OTUs and NTUs determined by both methods can be found in **Dataset S2 and S6**. Beta diversity analysis was performed on three taxonomic marker genes, *rplP*, *rplB*, and 16S rRNA NTUs. To ensure consistent and objective observations, both unrarefied and rarefied data (250 for *rplP* and *rplB*; 2000 for 16S rRNA NTUs) were analysed. Bray-Curtis dissimilarity was calculated and visualized using a non-metric multidimensional scaling ordination (NMDS) plot in phyloseq ^73^.

### Metagenomic assembly, binning, and phylogenomic analysis

Individual metagenomic assembly was performed with metaSPAdes v3.15.3 ^74^ with options min k: 27, max k: 127, and k step: 10. Bark metagenomes of *M. quinquenervia* were further co-assembled using MEGAHIT v1.2.9 ^75^ (min k: 27, max k: 127, and k step: 10). Contigs with lengths below 500 bp were discarded. Quality processed short reads were mapped to the assembled contigs using CoverM v0.6.1 (https://github.com/wwood/CoverM) with default parameters to generate contig coverage profiles. Genomic binning was performed using CONCOCT v1.1.0 ^76^, MaxBin2 v2.2.7 ^77^, and MetaBAT2 v2.15 ^78^ on *M. quinquenervia* bark metagenome contigs with length over 2000 bp. Resulting bins from the same assembly were then dereplicated using DAS_Tool v1.1.6 ^79^. Applying a common species-level threshold average nucleotide identity of 95%, bins from different assemblies were consolidated to a non-redundant set of metagenome-assembled genomes (MAGs) using dRep v3.4.3 ^80^. Only MAGs with a minimum completeness of 50% and a maximum contamination of 10%, as assessed by CheckM2 v1.0.2 ^81^, were retained. In total, 31 high-(completeness > 90% and contamination < 5%) and 83 medium-quality (completeness > 50% and contamination < 10%) ^82^ MAGs were recovered. Their corresponding taxonomy was assigned by GTDB-Tk v2.3.2 ^83^ with reference to the Genome Taxonomy Database (GTDB) R08-RS214 ^84^. All 114 MAGs were not classified to known species.

Phylogenomic trees were constructed to visualize the phylogenetic breadth of the 114 bacterial MAGs using the GTDB method ^85^. A concatenated multiple protein sequence alignment was built using the 120 GTDB bacterial marker genes in all bacterial MAGs by GTDB-Tk v2.3.2. IQ-TREE v2.2.0 ^86^ was then used to construct a maximum likelihood phylogenetic tree with 1000 ultrafast bootstrap replicates ^87^ under the WAG model of protein evolution with gamma-distributed rate heterogeneity (WAG + G20). The tree was rooted between Gracilicutes and Terrabacteria according to Coleman and Davin *et al.* ^88^.

### Metabolic annotation and genome analysis

To holistically estimate the metabolic capability of the bark communities to use trace gases and VOCs, metagenomes were searched against a custom protein database (https://github.com/GreeningLab/GreeningLab-database/tree/main) ^89^. The representative metabolic markers include 31 proteins involved in methane cycling (MmoX, PmoA, McrA, rMcrA), hydrogen cycling (large subunit of [NiFe]-, [FeFe]-, [Fe]-hydrogenases), carbon monoxide oxidation (CoxL, CooS), alkane and alkene oxidation (AcrA, HmoA, PxmA, BmoX, EtnC, PrmA, ZmoA), volatile aromatic compound oxidation (TmoA, IsoA, DmpN), nitrogen cycling (NosZ, NorB, Nod, NarG, NapA, NirS, NirK, NrfA), and aerobic respiration (CoxA, CcoN, CyoA, CydA). We further expanded the database with representative marker genes involved in VOC metabolism (methanol, acetone, alkane) and fermentation (ethanol, lactate, pyruvate, acetate, formate) from the KEGG prokaryotic protein database (accessed 22 November 2021) ^90^. Descriptions and annotations of genes are summarized in **Dataset S3**. To allow fair comparison of metagenomes from various sources, only quality-filtered forward reads longer than 130 bp by SeqKit v2.0.0 ^91^ were used for the analysis. Reads mapped to the 47 publicly available tree species genomes (*Melaleuca*, *Casuarina*, *Avicennia*, *Acacia*, *Persia*, *Macadamia*, *Telopea*, and *Eucalyptus*; genome information detailed in **Dataset S7**), were further filtered using bbmap.sh (options: maxindel=3 minid=0.7 minhits=2 bwr=0.16 bw=12 untrim quickmatch fast) implemented in BBTools v39.01 to reduce non-specific host read mapping to the metabolic database.

Metagenomic reads were searched against the gene databases using DIAMOND v.2.1.6 (query cover > 80%) ^92^. Results were filtered based on an identity threshold of 50%, except for CoxL, MmoX, BmoX, TmoA, IsoA, DmpN, EtnC, PrmA, ZmoA, [FeFe] and group 4 [NiFe]-hydrogenases (all 60%) ^89^. Subgroup and family classification of reads was based on the closest match to the sequences in databases while hydrogenase sequences were classified based on HydDB ^55^. Read counts for each gene were normalized to reads per kilobase per million (RPKM) by dividing the actual read count by the total number of reads (in millions) and gene length (in kilobases). Reads were also screened for the 14 universal microbial single copy ribosomal marker genes used in SingleM v.0.13.2 and PhyloSift ^93^ by DIAMOND (query cover > 80%, bitscore > 40) and normalized as above. The average gene copy number of a gene in the community was then estimated by dividing RPKM value of the gene by the geomean of RPKM value of the 14 universal single copy ribosomal marker genes. DRAM v1.2.4 ^94^ was used to perform metabolic annotations of MAGs with the Carbohydrate-Active enZYmes (CAZy) in dbCAN2 database v10 ^95^ and KEGG protein database. In addition, all MAGs and unbinned metagenomic assemblies were also screened for the presence of metabolic marker genes in our custom database. MAG and unbinned contig annotation results can be found in **Dataset S4 and S5**.

### Phylogenetic analysis of key trace gas metabolising genes

Maximum-likelihood phylogenetic trees were constructed to appraise the diversity and distribution of bark-associated microbes metabolizing major trace gases. Protein sequences of the catalytic subunits of [NiFe]-hydrogenase, [FeFe]-hydrogenase, CO dehydrogenase (CoxL), methyl-CoM reductase (McrA), particulate CH_4_ monooxygenase (PmoA), and soluble diiron monooxygenase family (MmoX, BmoX, TmoA, DmpN, EtnC, PrmA) from MAGs and unbinned contigs were first clustered and dereplicated at a 97% identity threshold using MMSeqs2 release 13-45111 (query cover > 80%, sensitivity: 7) ^96^. Only sequences with at least 200 amino acid residues or 40% query coverage with reference sequences in our database were used and aligned using MAFFT-L-INS-i v7.505 90 ^97^. Best-fit substitution model for each gene was determined using ModelFinder ^98^ implemented in IQ-TREE2 v2.2.0 ^86^ ([NiFe]-hydrogenase: LG+F+I+I+R7; [FeFe]-hydrogenase: Q.pfam+I+I+R5; CoxL: LG+F+R5; McrA: LG+I+G4; PmoA: LG+G4; soluble diiron monooxygenase: Q.pfam+I+I+R5). Maximum likelihood phylogenetic trees were constructed using IQ-TREE2 v2.2.0 ^86^ with 1000 ultrafast bootstrap replicates with the option -bnni to reduce impact of model violations ^87^. All trees were rooted at mid-point and visualized using iTOL v6 ^99^.

### Trace gas consumption and production experiments

Microcosms using *M. quinquenervia* and *C. glauca* bark samples were set up to determine the capacity of microbial communities to consume H_2_, CO, and CH_4_ aerobically and produce these trace gases anaerobically. For each sample in biological triplicate, 3 g of bark was cut into strips, transferred to a sterile 120-ml serum vial, and incubated at 20°C in the dark. For the oxic condition, vials were crimped sealed with butyl rubber stoppers (Sigma-Aldrich) and ambient air headspace was amended with an initial mixing ratio of approximately 10 parts per million (ppm) each of H_2_, CO, and CH_4_ (via a mixed gas cylinder containing 0.1 % v/v H_2_, CO, and CH_4_ each in N_2_, BOC Australia). For the anoxic condition, vials were crimped sealed with thick black rubber stoppers (to reduce O_2_ contamination during sampling; Rubberbv) and headspace was flushed with ultra-pure N_2_ (99.999%; BOC Australia) for 2 min. All incubations were performed at 20°C in the dark and all stoppers used were pre-treated with 0.1 N hot NaOH solution according to the description by Nauer *et al*. ^100^ to reduce abiotic emissions of H_2_ and CO from the stopper. Headspace H_2_, CO, and CH_4_ concentrations were monitored using a VICI gas chromatographic machine with a pulsed discharge helium ionization detector (model TGA-6791-W-4U-2, Valco Instruments Company Inc.) and an autosampler as previously described ^101^. The machine was regularly calibrated against ultra-pure H_2_, CO, and CH_4_ standards down to the limit of quantification (H_2_: 20 ppb; CO: 9 ppb; CH_4_: 500 ppb). For each condition, an empty vial was included as a negative control.

### Tree stem trace gas flux measurement

*In situ* tree stem gas fluxes were collected by first attaching a 30 cm × 30 cm semi-rigid chamber to each tree stem as described by Siegenthaler *et al*. ^102^. CO_2_ and CH_4_ fluxes were first measured by connecting the chamber to a portable gas analyser (CRDS, G4301 Gas Scouter, Picarro) via gas tubing (Bev-A-Line IV) passing through a drying desiccant (Drierite) within a closed loop. The tree stem fluxes were measured from between 40 - 70 cm above the soil or water level. On trees featuring uneven bark surfaces, occasionally, white potting clay was used to fill any gaps between the chambers and tree stems. Once chambers were sealed onto the tree stems, CH_4_ and CO_2_ concentration was recorded on the CRDS until a clear linear flux rate was observed for 150 s on high CH_4_ fluxing trees, and up to 15 min on low CH_4_ fluxing trees. The CRDS was factory-calibrated and features a precision of ± 0.3 ppb and lower detection limit of 0.9 ppb. *In situ* tree stem trace gas fluxes of H_2_, CO, and CH_4_ were then collected using the same chamber (as above) but with the inlet and outlet ports sealed with 2-way luer locks. Immediately after attaching each chamber, a 15 ml sample of the headspace was collected using a syringe every five minutes between time 0 to 20 min (n = 5). Each sample was injected into pre-evacuated 12ml Exetainer vial modified according to Nauer *et al*. ^100^ to optimise sample storage and avoid CO contamination. The change in H_2_, CO, and CH_4_ over each time step was analysed on the gas chromatograph (described above). Duplicate atmospheric trace gas samples were collected at each forest site. The trace gas fluxes (*F*) for trees were calculated using the equation:

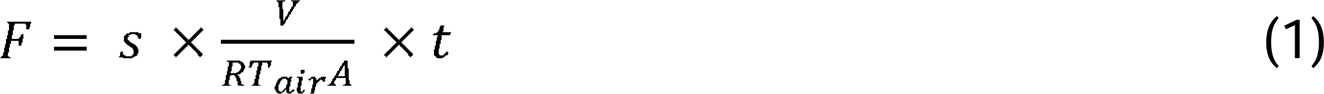

where *s* is the regression slope for each chamber incubation deployment (ppm sec^−1^), *V* is the chamber volume (m^3^), *R* is the universal gas constant (8.205 × 10^−5^ m^3^ atm^−1^ K^−1^ mol^−1^), *T*_air_ is the mean air temperature in Kelvin (K), *A* is the surface area of each chamber (m^2^), and *t* is the conversion factor from second to day, and to mmol of CH_4_. The *s* and *T*_air_ terms were extracted using a modified “*GasFlux*” package (https://git-dmz.thuenen.de/fuss/gasfluxes). The *V* term within CRDS flux calculations (i.e., total volume of the closed loop including chamber, 4 m length of gas tubing, desiccant canister and internal volume of CRDS) was calculated as described by Jeffrey *et al*. 2020 ^103^.

### Sapwood stem-gas composition

Sapwood gas was collected as described in Jeffrey *et al*., 2024 ^22^. Briefly a 5 × 5 cm swatch of bark was removed using a sterile blade to reveal sapwood surface. A 12 mm diameter hole, 11 cm deep was drilled into the sapwood to create a cavity of ∼12 ml volume. A rubber stopper was inserted into the cavity opening and left for the internal stem gas composition of composition of H_2_, CO and CH_4_ to equilibrate for at least 4 h. Using two 20 ml syringes, a 12 ml of ambient air sample was mixed into the cavity whilst carefully extracting stem-equilibrated gas into the second empty syringe. The process was carefully repeated four times to mix the ambient air with stem gas sample. The 12 ml sapwood gas sample was then injected into a pre-evacuated Exetainer modified for trace gases, and analysed on the gas chromatograph (as described above). The 1:1 dilution with atmospheric air was back-calculated to determine original sapwood stem-gas concentrations of each tree species.

### Quantification and identification of bark-emitted volatile organic compounds (VOCs)

The concentration of VOCs within the bark of a flooded *M. quinquenervia* tree at two stem heights (40 and 145 cm) was determined using a Proton Transfer Reaction Time of Flight Mass Spectrometer (PTR-ToF-MS, Ionicon Analytik). First, to collect 1 L of VOC gas sample, an air-tight semi-rigid chamber (as described above) was attached to the tree at the specified heights. The chamber outlet was then connected in-line via gas tubing (Bev-A-Line IV) to a desiccant trap (Drierite), a micro diaphragm pump (Parker, T2-05) and a 1 L dual sampling port - foil gas bag (custom made, Cali-5-Bond), that returned to the stem chamber inlet, creating a closed loop. The stem chamber headspace was gently circulated within the closed loop using the air pump at 30% flow rate (0.24 L/minute) for 24 h to allow for the chamber and closed loop to equilibrate with the tree stem gases and VOCs. Duplicate 5 mL subsamples from the stem gas bags were analysed for stem CH_4_ and CO_2_ concentrations using a small sample induction module (SSIM AO314) connected to a CRDS (Picarro, G2201-i).

Following sample collection, the 1 L gas bags were analysed in the lab using the PTR-ToF-MS ^104^. The instrument was calibrated using two certified gas standards (Apel Riemer Environmental) containing a total of 20 VOC species including methanethiol, acetone, dimethyl sulfide, isoprene, 1,8-Cineole and methyl iodide (gas mix D024451) and methanol, acetonitrile, acetaldehyde, acetone, methacrolein, methyl ethyl ketone, benzene, toluene, xylene, chlorobenzene, 1,3,5-trimethylbenzene, α-pinene, 1,2-dichlorobenzene and 1,2,4-trichlorobenzene (gas mix D024442) at ∼ 1 ppm each. The two gas mixtures (D024451 and D024442) as well as blanks of ultra-high purity N_2_ were consecutively connected to the instrument via a liquid calibration unit (Ionicon Analytik) at flow rates of 10 standard cubic cm per minute (sccm) and 5 sccm, respectively, with an ultra-high purity N_2_ carrier gas at a flow rate of 1000 sccm. Following calibration, samples were consecutively processed by simply connecting the Cali-5-Bond gas bags to the instrument’s inlet until the bags were nearly empty (10-15 min). A Cali-5-Bond gas bag previously flushed 3 times and filled with ultra-high purity N_2_ was also measured as a blank.

VOCs identification and quantification were performed using the IONICON PTR-MS Viewer v.3.4.5 software. Gaussian functions were employed to satisfactory fit mass spectrometric peaks, thus reducing the computational effort at the cost of introducing a resolution error 20% ^105^. A mass calibration was applied across all spectra to ensure accurate mass measurements throughout the analysis. Total VOC concentrations were estimated by compiling all the measured VOCs based on their nominal masses within the analytical mass range (0 to 640 m/z). Correction factors were calculated based on the known VOC concentrations in the certified gas mixes and applied to the estimated VOC concentrations obtained via the PTR-MS Viewer. Peaks of interest in each sample (i.e. at 40 and 145 cm stem heights) were identified by comparing the blank-corrected sample spectra with each other using a factor of 5x. Each peak of interest was defined using the inbuilt NIST library and compounds of interest were identified based on their level of correctness.

### Trace gas flux calculation and modelling

Results from the two microcosm experiments were converted into gravimetric flux rates (flux/g of bark/d) and areal flux rates (flux/m^2^ of tree stem/d reported as µmol m^-2^ d^-1^). In the oxic treatments, the three gases were consumed at different rates so the periods of most linear decrease were used to calculate oxidation rates. Therefore, the time steps between 0 – 4 h, 2 – 21 h, and 2 – 94 h were used for H_2_, CH_4_, and CO, respectively. For the anaerobic treatment, production of H_2_, CH_4_, and CO was much slower, therefore the time steps between 0 – 85 h were used to estimate gas production rates. For the axial bark microcosm experiment (collected from multiple stem heights up to 9 m), for the aerobic treatment, the time steps of 0 – 6 h was used for both CH_4_ and H_2_, and the 6 – 48 h time step for CO. For the axial bark anaerobic incubations, the 0 – 48 h for CO and CH_4_, and 0 – 168 h for H_2_ time steps were used. Bottle concentrations were converted to µmol (at standard temperature and pressure) before conversion to bark gravimetric (nmol) and areal flux rates (µmol). To compare soil trace gas flux literature to our tree stem H_2_, CO and CH_4_ fluxes, we upscaled to the ecosystem level (i.e., accounting for tree size and forest density), using the mean *in situ* stem flux rates of each gas (i.e., µmol m^-2^ d^-1^ as per above). These were scaled only to 6 m of stem height (halfway to canopy) and multiplied by the mean surface area of bark/hectare, as well characterised previously by Jeffrey *et al*. 2023 ^16^.

## Footnotes

### Data Availability

All raw metagenomes, metagenomic assemblies, and metagenome-assembled genomes of the samples were deposited to the National Center for Biotechnology Information (NCBI) Sequence Read Archive. Note that deposited data will be made publicly available upon manuscript acceptance. All other study data are included in the article and/or supporting information.

## Acknowledgements

This study was supported by research grants from Australian Research Council (DE240100338 to L.C.J; DP210100096 to D.T.M, S.G.J; DE230101346 to S.K.B; LP160100061 to S.G.J.; LE200100155 to E.D.), SRIEAS grant Securing Antarctica’s Environmental Future (SR200100005; salary support to S.K.B), the Hermon Slade Foundation (to L.C.J., D.T.M., S.G.J, C.G.), Australian Institute of Nuclear Science and Engineering Postgraduate Research Award (to L.C.J), a Monash Summer Studentship (to M.H.), a Holsworth Wildlife Research Endowment fund (to P.M.L.), a Monash FMNHS Early Career Postdoctoral Fellowship (ECPF23-1113137961; to P.M.L.), and an NHMRC EL2 Fellowship (APP1178715; to C.G.). We greatly appreciate the MonARCH HPC Cluster and the MASSIVE HPC facility for providing computation platforms and resources. We also thank Dr. Sebastian Euler, Dr. Mitchell Call, Dr. Charly Moras, Scott Cramb, Evagzard Kipnusu and Billie Hopkins for fieldwork assistance, and Dr. Elenora Chiri and Dr. Philipp Nauer for helpful discussion.

## Author contributions

L.C.J., P.M.L., C.G., and D.T.M. conceived, designed, and supervised the study. Different authors were responsible for field sampling (L.C.J., J.D., P.G.A., S.G.J.), greenhouse gas flux measurement (L.C.J., J.D.), *in situ* stem gas sampling (J.D., L.C.J.), bark microcosm experiments (M.H., P.M.L., J.D., P.G.A., D.T.M., L.C.J.), gas chromatography (M.H., T.J., T.H., P.M.L.), volatile organic compound collection and analysis (P.G.A., E.D., J.D., L.C.J.), community DNA extraction and quantitative PCR assay (T.J.), metagenomic sequencing, assembly and binning (P.M.L., S.K.B.), genome analysis (P.M.L.), metabolic annotation (P.M.L., S.K.B., C.G., X.D., N.V.C.), community analysis (P.M.L., S.K.B., P.G.A.), and phylogenetic analysis (P.M.L.). P.M.L. and L.C.J. analysed data and wrote the manuscript with input from all authors. All authors edited and reviewed the manuscript.

