## Supporting Information for "Wetland tree barks are dynamic hotspots for microbial trace gas cycling"

**Supplementary Table**

**Table S1**. Stem gas composition collected from eight *M. quinquenervia*.

| Tree ID | $H_2$ | $CH_4$ (ppm) | CO |
|---|---|---|---|
| 1 | 1.31 | 3378 | 0.77 |
| 2 | 14.13 | 223.9 | 6.97 |
| 3 | 13.84 | 159.3 | 13.98 |
| 4 | 3.23 | 135.7 | 6.11 |
| 5 | 1.53 | 367.0 | 1.25 |
| 6 | 14.46 | 511.5 | 8.68 |
| 7 | 1.17 | 2629 | 1.04 |
| 8 | 1.52 | 1241 | 0.66 |
| Average | 6.40 | 1081 | 4.93 |
| Min | 1.17 | 135.7 | 0.66 |
| Max | 14.46 | 3378 | 13.98 |

**Table S2.** Summary of major VOC composition from lower and upper tree stems of *M. quinquenervia*.

| VOC | Lower stem (ppb) | Upper stem (ppb) | mass |
|---|---|---|---|
| Acetaldehyde | 261.75 | 207.65 | 33 |
| Acetone | 130.89 | 120.87 | 34 |
| Butanal | 110.05 | 111.76 | 41 |
| Methanol, methyl alcohol | 104.38 | 75.08 | 43 |
| Propene | 83.96 | 540.72 | 44 |
| Cyclopropene, Propyne, Methylacetylene | 81.07 | 391.64 | 45 |
| Butene | 71.33 | 50.95 | 46 |
| Formic acid | 57.16 | 13.47 | 47 |
| Hydroxylamine | 13.56 | 14.47 | 57 |
| Propanamide | 6.56 | 6.71 | 59 |
| Formamide, Methane, nitroso- | 6.53 | 5.39 | 63 |
| Propylene Glycol, 2-methoxy-ethanol, Propanediol | 4.84 | 3.36 | 70 |
| 2,5-Piperazinedione, N-α-Acetylglycinamide,Ethyl diazoacetate, Dimethylfurazan monoxide | 4.59 | 0.85 | 73 |
| Cyclopropane, 1,1-diethynyl- | 4.27 | 7.75 | 74 |
| Isopropyl diazoacetate | 3.60 | 1.13 | 75 |
| Acetaldimine, Ethylenimine | 3.35 | 20.97 | 77 |
| Cyclopentyl radical | 2.72 | 2.38 | 91 |
| Propanoic acid | 2.14 | 2.02 | 101 |
| Dimethyl sulfide | 1.78 | 1.28 | 115 |
| Hexanal | 0.92 | 2.09 | 129 |

**Table S3.** Summary of the multiple field campaigns, tree species, sampling purpose, ancillary parameters and site locations used for this study.

| Date | Purpose | Species | Qty | Sample height (cm) | DBH (cm) | Water depth (cm) | Sample description | Latitude | Longitude | Bark VWC (%) |
|---|---|---|---|---|---|---|---|---|---|---|
| 20/05/2020 | qPCR and metagenomic sequencing | *M. quinquenervia* | 8 | - | - | - | 8 bark, 2 soil and 2 water samples | -28.351481° | 153.571267° | - |
| 10/05/2021 | Metagenomic sequencing | *A. marina* | 1 | *84* | *13.5* | - | 8 samples from various species (moist conditions) | -28.357869° | 153.572874° | 21% |
| | | *E. propinqua* | 1 | 85 | 50.8 | - | | -28.368982° | 153.563713° | 21% |
| | | *E. siderophloia* | 1 | 84.5 | 43.8 | - | | -28.369051° | 153.563095° | 9% |
| | | *B. intergrofolia* | 1 | *84.5* | 43.4 | - | | -28.351384° | 153.572168° | 10% |
| | | *A. longifolia* | 1 | 105 | 14.8 | - | | -28.349347° | 153.571034° | 12% |
| | | *C. glauca* | 1 | 35 | 31.5 | 63 | | -28.351939° | 153.571691° | 37% |
| | | *L. suaveolens* | 1 | 35 | 29.4 | 23 | | -28.349192° | 153.571007° | 33% |
| | | *M. quinquenervia* | 1 | 61 | 23 | 29 | | -28.349304° | 153.571015° | 63% |
| 3/11/2021 | Trial microcosm experiment | *M. quinquenervia* | 1 | 20-50 | - | - | Bark sample | -28.350509° | 153.571657° | - |
| | | *C. glauca* | 1 | 20-50 | - | - | | -28.351910° | 153.571664° | - |
| 21/01/2022 | Microcosm experiment | *M. quinquenervia* | 1 | 20-50 | - | - | Bark sample | -28.350509° | 153.571657° | - |
| | | *C. glauca* | 1 | 20-50 | - | - | | -28.351910° | 153.571664° | - |
| 31/03/2022 | Bark VOC composition | *M. quinquenervia* | 1 | 40,145 | 26 | 90 cm | Flooded tree stem VOC sampled at two heights | -28.350289° | 153.571777° | - |
| 14/11/2023 | In situ trace gas fluxes | *M. quinquenervia* | 8 | 40-65 | 23-45 | 0 cm | 8 trees - stem fluxes and sapwood gas (dry) | -28.347774° | 153.570875° | - |
| 6/02/2024 | In situ trace gas fluxes | *M. quinquenervia* | 8 | 40-65 | 23-45 | 20 cm | 8 stem fluxes and sapwood gas (Wet) | -28.347774° | 153.570875° | - |
| 22/03/2024 | In situ trace gas fluxes and microcosm experiment | *M. quinquenervia* | 4 | 40-65 | 23-45 | 5 | Fluxes and bark axial stem locations to 8.8m | -28.347774° | 153.570875° | - |
| 20/05/2024 | Low affinity $CH_4$ oxidation assay | *M. quinquenervia* | 1 | 55 | 28 | 45 | 2 bark samples | -28.349304° | 153.571015° | 75% |

**Table S4.** Summary of triplicate *M. quinquenervia* and *C. glauca* bark microcosm assays showing aerobic consumption of $H_2$, CO and $CH_4$ trace gases (top) vs anaerobic production of $H_2$, CO and $CH_4$ trace gases (bottom). Values are normalised to the bark weight of each microcosm assay (nmol/g of bark/d) and upscaled to areal rate (µmol/m$^2$ of bark/d). Note: n.d indicates no detectable change in values.

| Aerobic treatment | Gas | Uptake (nmol/g of bark/d) | | | Areal uptake (µmol/m$^2$ bark/d) | | |
|---|---|---|---|---|---|---|---|
| *Melaleuca* bark | $H_2$ | -46.4 | ± | 3.44 | -334.7 | ± | 24.80 |
| | CO | -5.8 | ± | 0.60 | -41.9 | ± | 4.33 |
| | $CH_4$ | n.d. | | | | | |
| *Casuarina* bark | $H_2$ | -73.9 | ± | 12.6 | -531.0 | ± | 90.44 |
| | CO | -3.1 | ± | 0.78 | -22.5 | ± | 5.60 |
| | $CH_4$ | n.d. | | | | | |

| Anaerobic treatment | | Production (nmol/g of bark/d) | | | Areal production (µmol/m$^2$ bark/d) | | |
|---|---|---|---|---|---|---|---|
| *Melaleuca* bark | $H_2$ | 42.6 | ± | 20.1 | 307 | ± | 145 |
| | CO | 7.1 | ± | 1.93 | 51.4 | ± | 13.9 |
| | $CH_4$ | 0.008 | ± | 0.05 | 0.061 | ± | 0.37 |
| *Casuarina* bark | $H_2$ | 70.1 | ± | 35.1 | 504 | ± | 252 |
| | CO | 0.23 | ± | 0.11 | 1.68 | ± | 0.81 |
| | $CH_4$ | 120 | ± | 117 | 862 | ± | 840 |

**Table S5.** Results and ancillary data of the high-concentration $CH_4$ oxidation microcosm experiment (~ 500 ppm $CH_4$) determined using a cavity ring-down spectrometer for two *M. quinquenervia* bark samples.

| Sample ID | Moisture content (%) | Bark weight (g) | Days | $CH_4$ oxidation rate (nmol/ g of bark/ day) | Areal $CH_4$ oxidation rate (µmol/ m$^2$ bark/ day) |
|---|---|---|---|---|---|
| *Melaleuca* 1 | 75% | 80.20 | 9.23 | -4.50 | -32.46 |
| *Melaleuca* 2 | 73% | 91.50 | 9.22 | -5.55 | -40.07 |
| Average | 74% | 85.85 | 9.23 | -5.02 | -36.26 |

**Table S6.** Summary of duplicate *M. quinquenervia* bark microcosm assays (collected from axial stem locations up to 8.8 m) showing aerobic consumption of $H_2$ and CO (top) vs anaerobic production of $H_2$, CO and $CH_4$ trace gases (bottom). Values are normalised to the bark weight of each microcosm assay (nmol/g of bark/d) and upscaled to areal rate (µmol/m$^2$ of bark/d). Note: n.d. indicates no detectable change in values for some $CH_4$ samples.

| Aerobic treatment | Gas | Uptake (nmol/g of bark/d) | | | Areal uptake (µmol/m$^2$ bark/d) | | |
|---|---|---|---|---|---|---|---|
| 0.4 m | $H_2$ | -74.08 | ± | 1.36 | -534.09 | ± | 9.83 |
| | CO | -4.12 | ± | 1.92 | -29.72 | ± | 13.81 |
| | $CH_4$ | n.d. | | | | | |
| 2.0 m | $H_2$ | -85.42 | ± | 2.67 | -615.83 | ± | 19.28 |
| | CO | -11.44 | ± | 0.69 | -82.45 | ± | 5.01 |
| | $CH_4$ | n.d. | | | | | |
| 6.0 m | $H_2$ | -85.79 | ± | 2.26 | -618.48 | ± | 16.31 |
| | CO | -10.65 | ± | 0.39 | -76.75 | ± | 2.82 |
| | $CH_4$ | n.d. | | | | | |
| 8.8 m | $H_2$ | -87.37 | ± | 0.49 | -629.88 | ± | 3.53 |
| | CO | -7.72 | ± | 0.20 | -55.67 | ± | 1.44 |
| | $CH_4$ | n.d. | | | | | |
| Anaerobic treatment | | | | | | | |
| 0.4 m | $H_2$ | 2.21 | ± | 1.02 | 15.96 | ± | 7.33 |
| | CO | 3.84 | ± | 0.91 | 27.72 | ± | 6.55 |
| | $CH_4$ | n.d | | | | | |
| 2.0 m | $H_2$ | 1.11 | ± | 0.13 | 7.99 | ± | 0.97 |
| | CO | 1.27 | ± | 0.14 | 9.19 | ± | 1.02 |
| | $CH_4$ | 192.53 | ± | 29.39 | 1388.03 | ± | 211.86 |
| 6.0 m | $H_2$ | 1.92 | ± | 0.69 | 13.81 | ± | 4.98 |
| | CO | 1.78 | ± | 0.06 | 12.84 | ± | 0.40 |
| | $CH_4$ | n.d | | | | | |
| 8.8 m | $H_2$ | 0.45 | ± | 0.05 | 3.24 | ± | 0.39 |
| | CO | 1.95 | ± | 0.46 | 14.09 | ± | 3.34 |
| | $CH_4$ | n.d | | | | | |

**Table S7**. Summary of *M. quinquenervia* tree stem trace gas flux rates of $CO_2$, $H_2$, $CH_4$, and CO between dry and wet *in situ* campaigns.

| Campaign | Tree | $CO_2$ flux | $H_2$ Flux | $CH_4$ flux | CO flux |
|---|---|---|---|---|---|
| | | | (µmol/m$^2$ bark/d) | | |
| Dry | 1 | 51.79 | -3.59 | 124.4 | 8.10 |
| | 2 | 73.42 | -3.63 | 34.26 | 5.58 |
| | 3 | 73.37 | -9.70 | 94.15 | 3.68 |
| | 4 | 42.18 | -7.16 | 66.08 | 4.93 |
| | 5 | 61.98 | -7.50 | 89.30 | 4.52 |
| | 6 | 195.2 | -3.24 | 132.3 | 12.21 |
| | 7 | 112.7 | -4.85 | 656.6 | 9.34 |
| | 8 | 33.40 | -6.66 | 45.22 | 10.57 |
| | Average | 80.50 | -5.79 | 155.3 | 7.37 |
| | SE ± | 18.49 | 0.82 | 72.7 | 1.11 |
| | Min | 33.40 | -9.70 | 34.3 | 3.68 |
| | Max | 195.2 | -3.24 | 656.6 | 12.21 |
| Wet | 1 | 238.0 | -8.23 | 22821 | 13.80 |
| | 2 | 91.90 | -18.33 | 3481 | 17.08 |
| | 3 | 146.7 | -11.81 | 3393 | 16.61 |
| | 4 | 21.31 | -13.79 | 3338 | 13.46 |
| | 5 | 94.59 | -15.27 | 4233 | 9.10 |
| | 6 | 135.8 | -29.36 | 8411 | 7.03 |
| | 7 | 173.7 | -13.60 | 20618 | 32.42 |
| | 8 | 88.35 | -13.92 | 15472 | 19.60 |
| | Average | 123.8 | -15.54 | 10221 | 16.14 |
| | SE ± | 23.08 | 2.22 | 2906 | 2.75 |
| | Min | 21.31 | -29.36 | 3338 | 7.03 |
| | Max | 238.0 | -8.23 | 22821 | 32.42 |

**Supplementary Figure**

**Figure S1. Summary of sampling location and tree species.** Images show differing bark and stems of multiple tree species used for microbial bark collection across the various forest biomes. The *in-situ* trace gas fluxes were performed within the freshwater wetland dominated by *M. quinquenervia* located to the north.

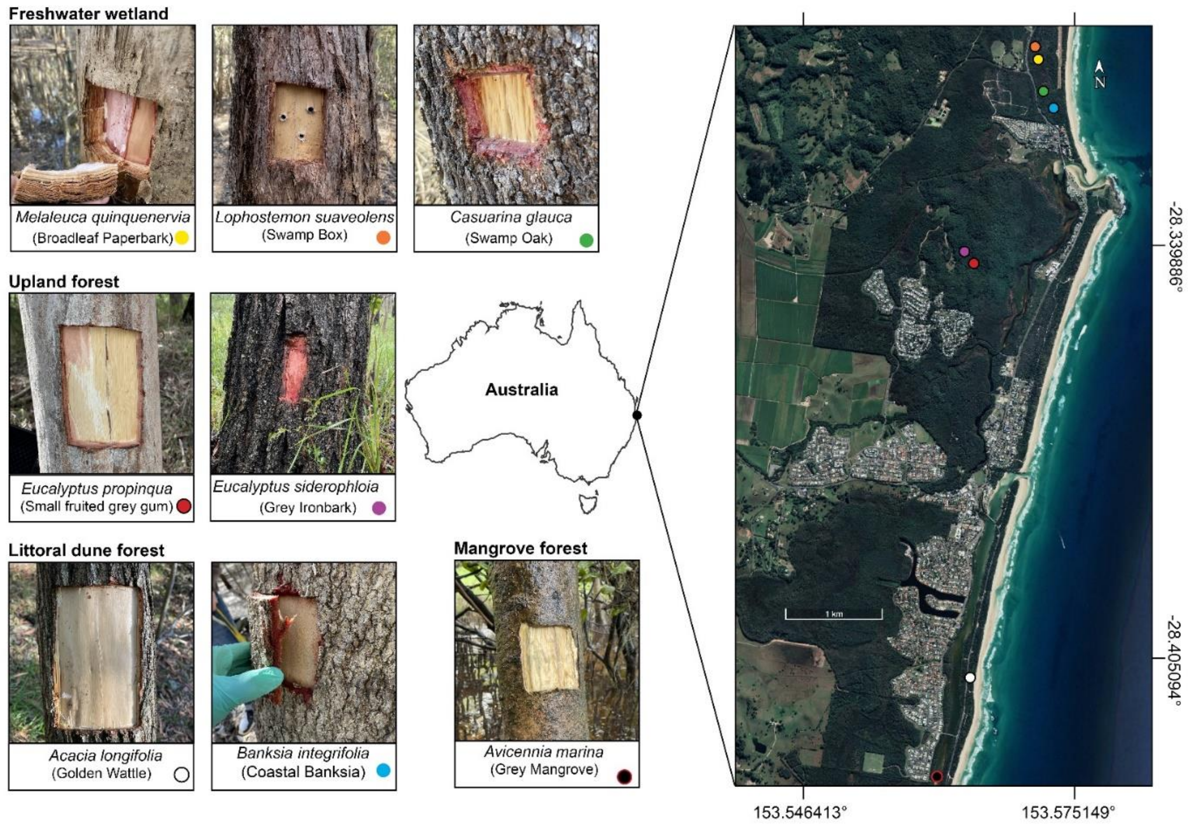

**Figure S2. Beta diversity analyses of the bark, soil, and water samples based on Bray-Curtis dissimilarity and visualized by a non-metric multidimensional scaling ordination (NMDS) plot.** Unrarefied and rarefied taxonomic marker genes were analysed. (**A**) ribosomal protein *rplP*. (**B**) ribosomal protein *rplB*. (**C**) 16S rRNA gene.

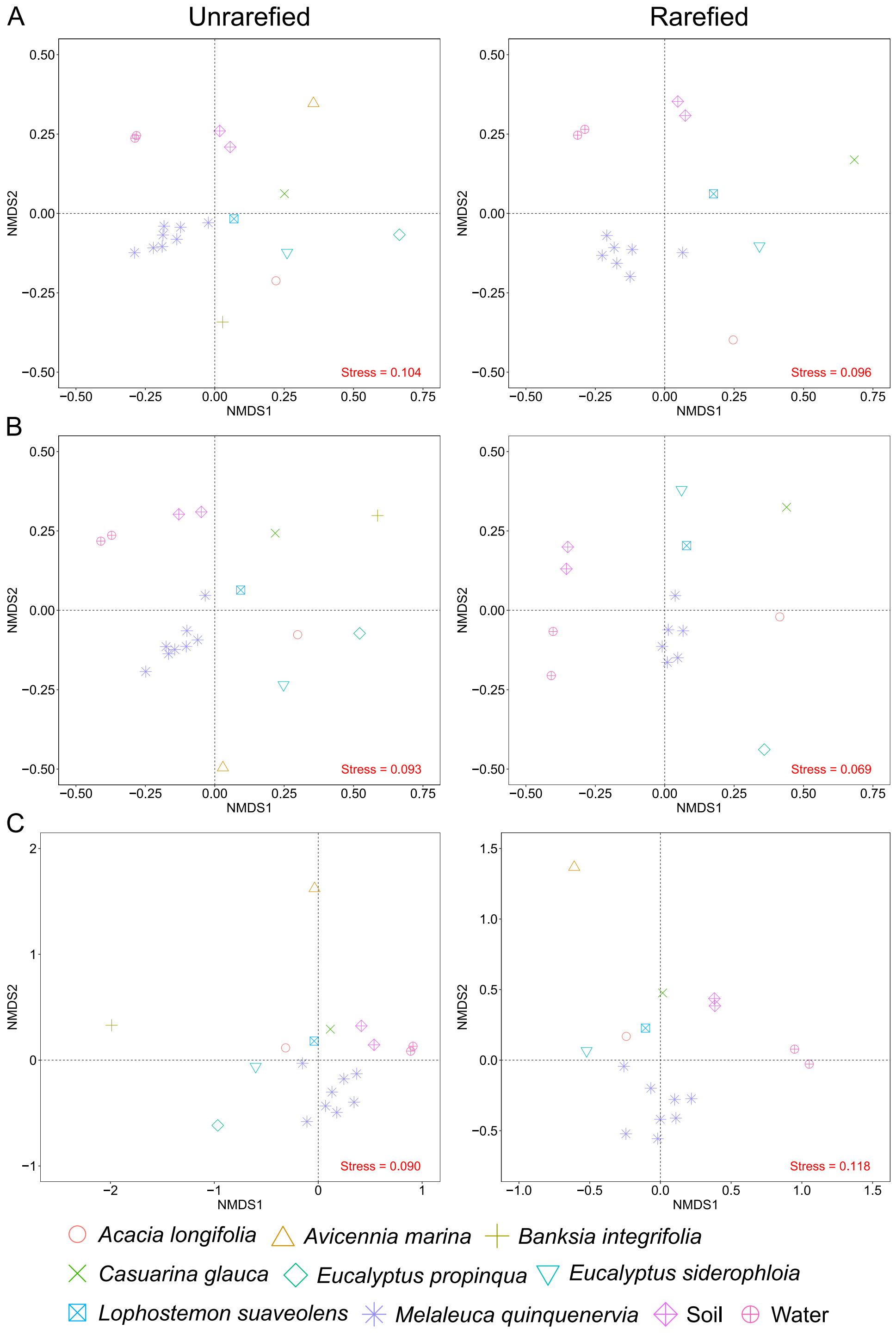

**Figure S3. Abundance and presence of metabolic marker genes involved in fermentation, volatile organic compound (VOC) degradation, denitrification, and carbohydrate degradation in bark, soil, and freshwater communities**. Heatmaps showing (**A**) the community average copy number of and (**B**) proportion of MAGs encoding metabolic marker genes for fermentation (amino acid degradation, pyruvate decarboxylation, formate / acetate / ethanol / L-lactate / D-lactate fermentation), VOC metabolism (toluene / isoprene / alkane / hydrocarbon oxidation), and denitrification (nitrate / nitrite reduction). (**C**) Barchart showing the average copy number of the six families of carbohydrate-active enzymes (CAZyme) normalized by genome size (per megabase) in metagenome-assembled genomes (MAGs).

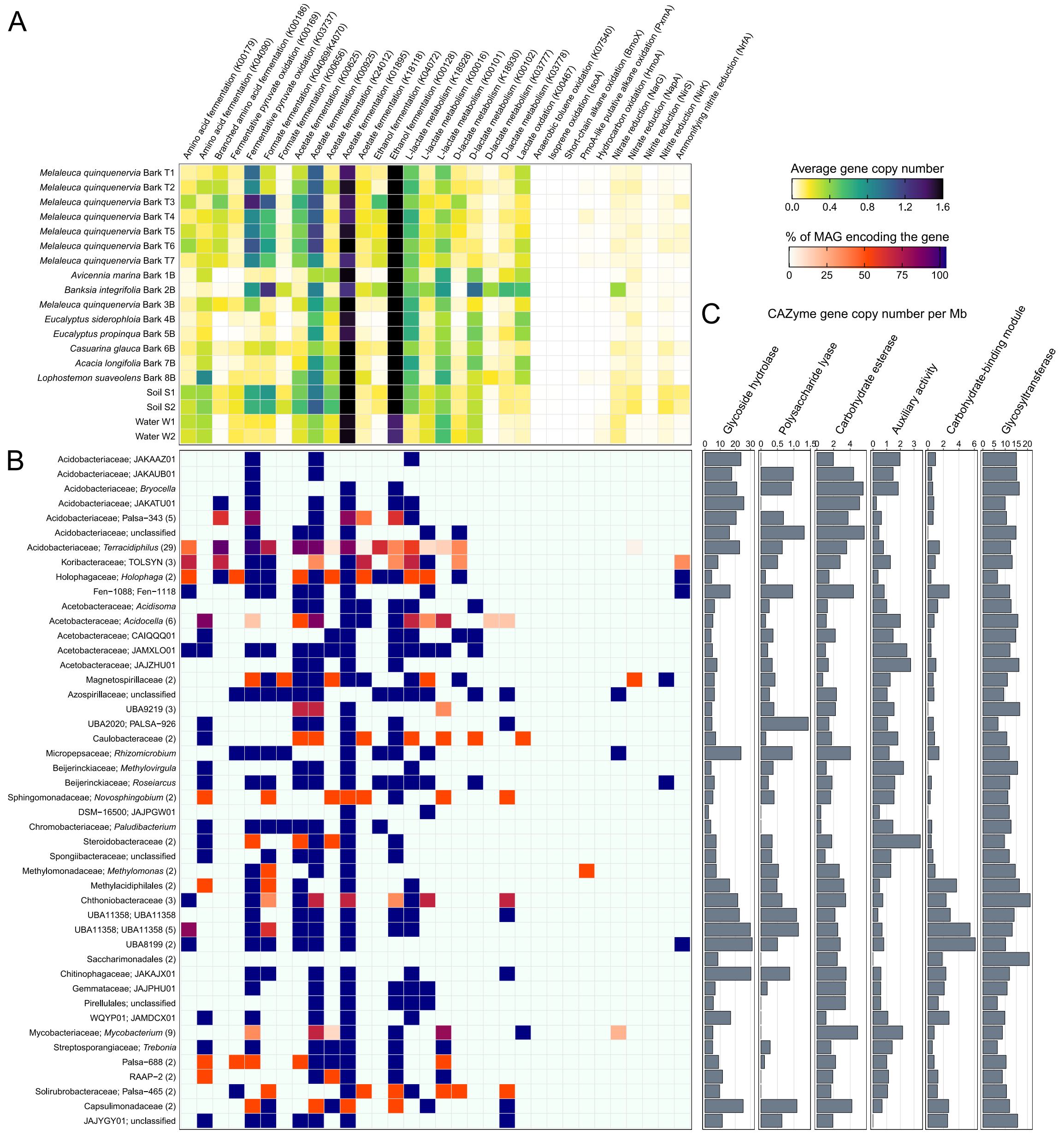

**Figure S4. Maximum-likelihood phylogenetic tree of [NiFe]-hydrogenase large subunits from both MAGs and unbinned contigs dereplicated at 97% identity**. Trees were rooted at mid-point, nodes with >75% branch support (1000 ultrafast bootstrap replicates) were shown in black dots, and the scale bar indicates the average number of substitutions per site. Hydrogenase subgroups and functions were classified based on the hydrogenase database (HydDB) [1]. The colored leaf nodes denote phylum-level taxonomy of MAGs and environmental origins of unbinned hydrogenase sequences.

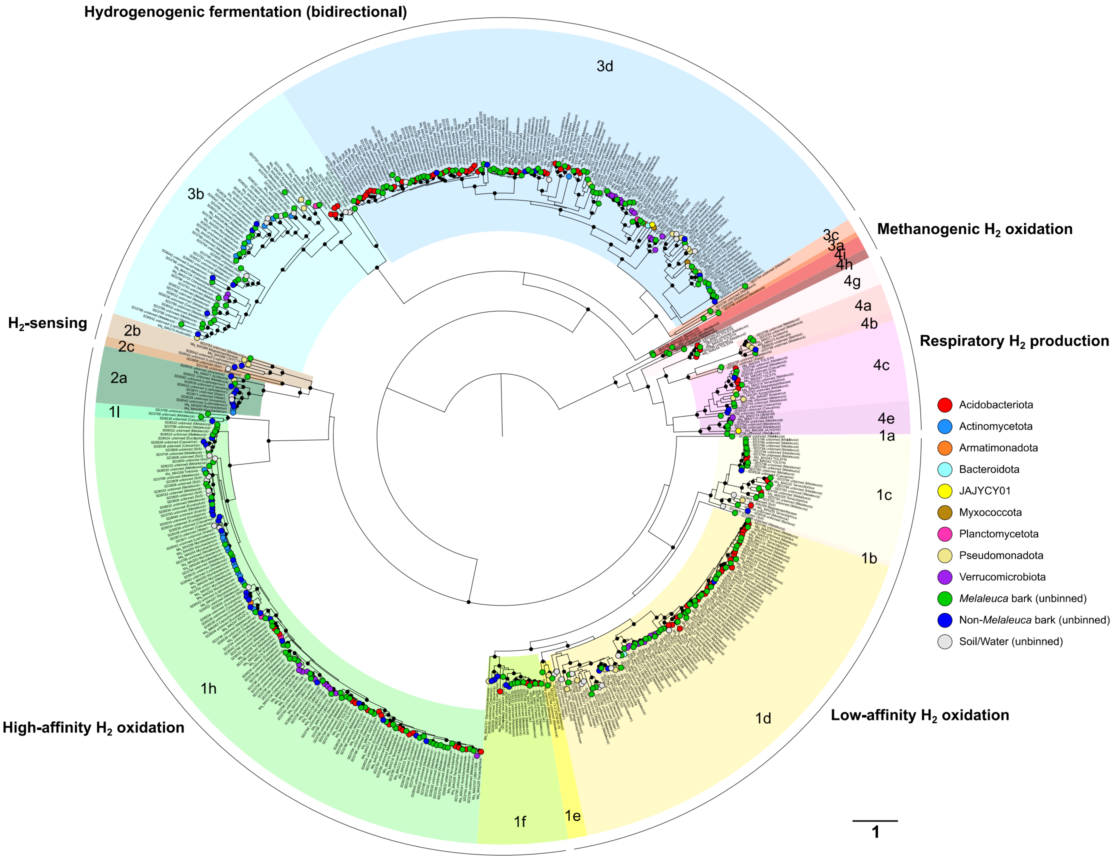

**Figure S5**. **Maximum-likelihood phylogenetic tree of carbon monoxide dehydrogenase catalytic subunits (CoxL) from reference sequences, MAGs, and unbinned contigs dereplicated at 97% identity**. Trees were rooted at mid-point, nodes with >75% branch support (1000 ultrafast bootstrap replicates) were shown in black dots, and the scale bar indicates the average number of substitutions per site. CoxL clades were classified based on ref [2]. The colored leaf nodes denote phylum-level taxonomy of MAGs and environmental origins of unbinned sequences.

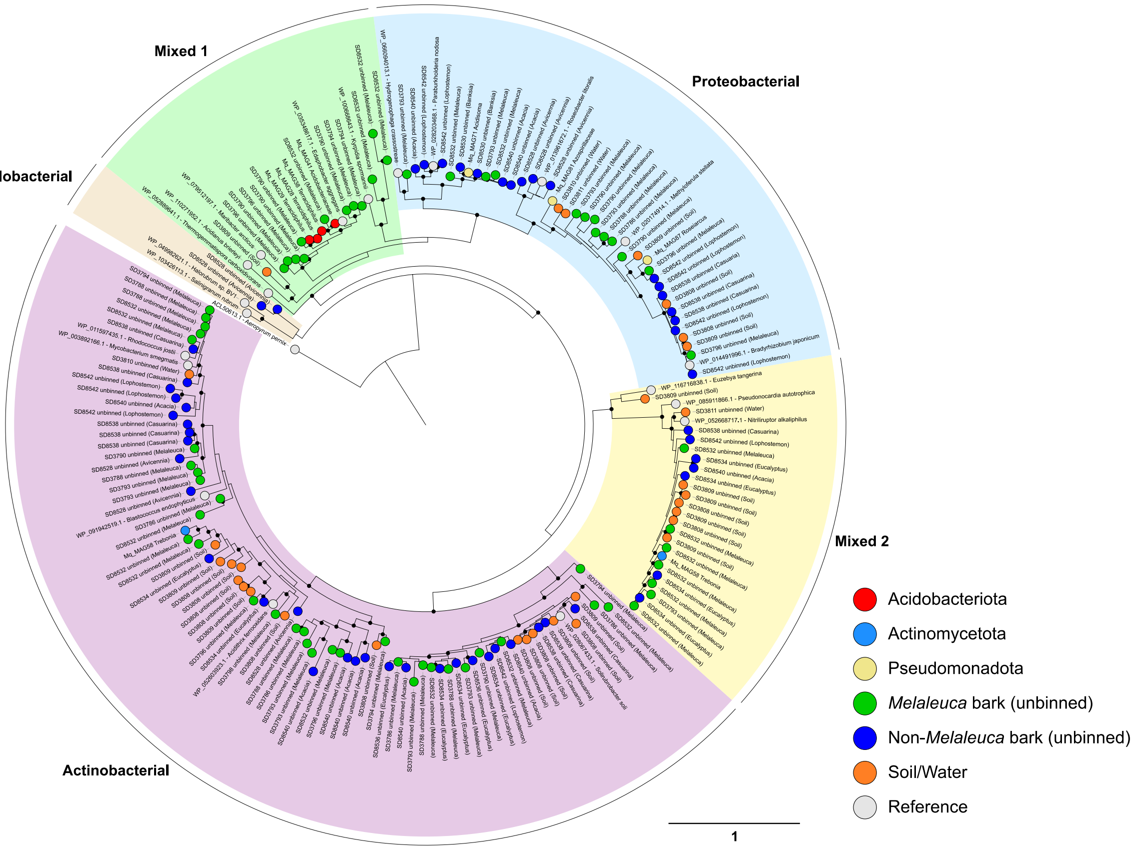

**Figure S6**. **First campaign of gas chromatography measurements of bark-mediated hydrogen, carbon monoxide, and methane metabolism under oxic and anoxic conditions**. Microcosms of *M. quinquenervia* T4 (red line) and *C. glauca* B6 (blue line) barks (n = 3 technical replicates) collected on 3rd Nov 2021 were incubated at 20°C, with empty vials as negative controls (black line). Oxic incubation was set up with approximately 10 ppmv each of $H_2$, CO, and $CH_4$ in the ambient air headspace while anoxic incubation was set up by purging headspace with ultra-pure nitrogen gas. Data are presented as mean ± S.D. values of headspace (**A**) $H_2$, (**B**) CO, (**C**) $CH_4$ mixing ratios.

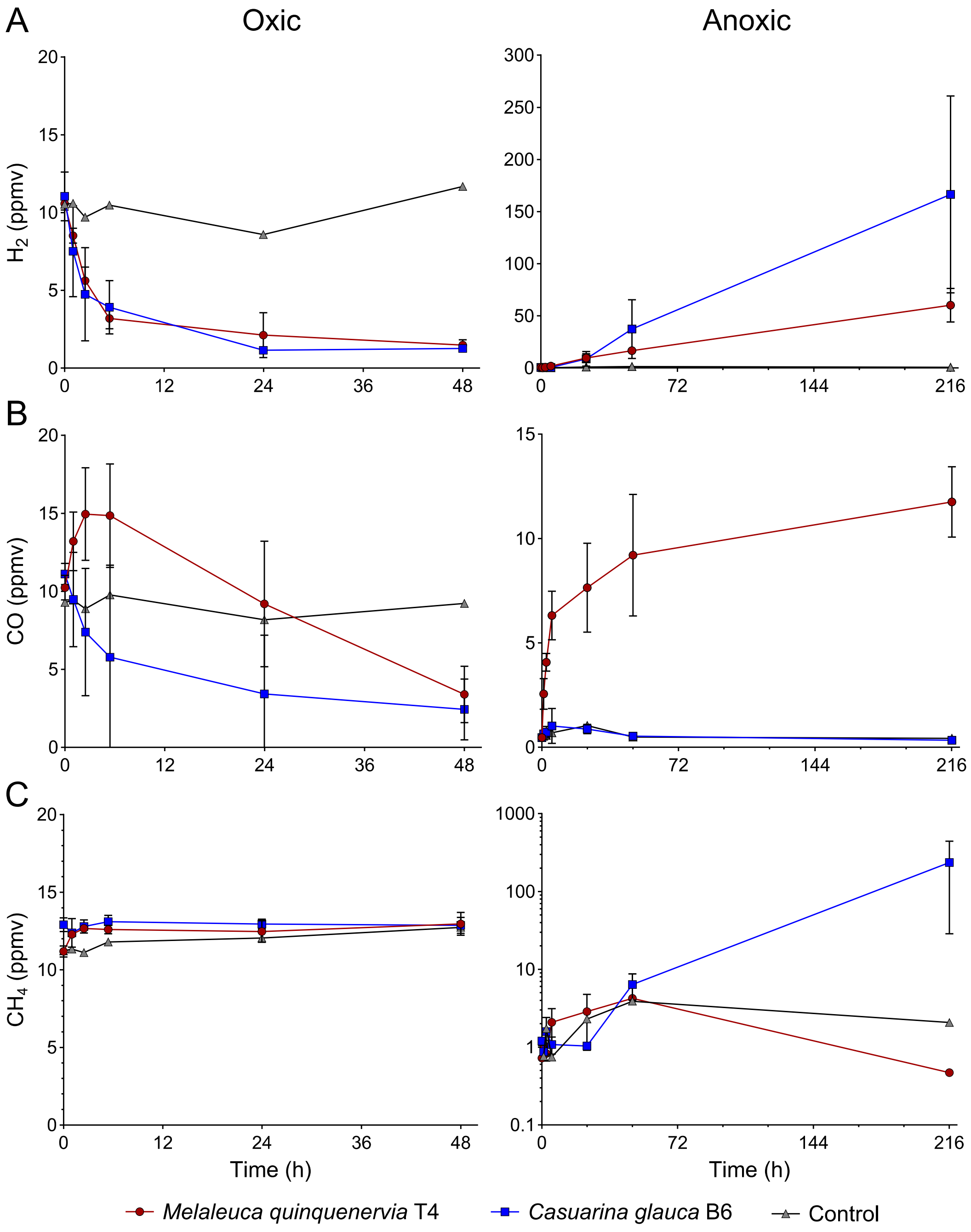

**Figure S7. Microcosm results** of aerobic (top) vs anaerobic (bottom) *M. quinquenervia* bark assays of bark collected from various axial stem heights and measured in duplicate, showing the potential of active trace gas production and consumption within caulosphere microbial communities. Values of each microcosm assay are normalised to the bark weight (nmol/g of bark/d).

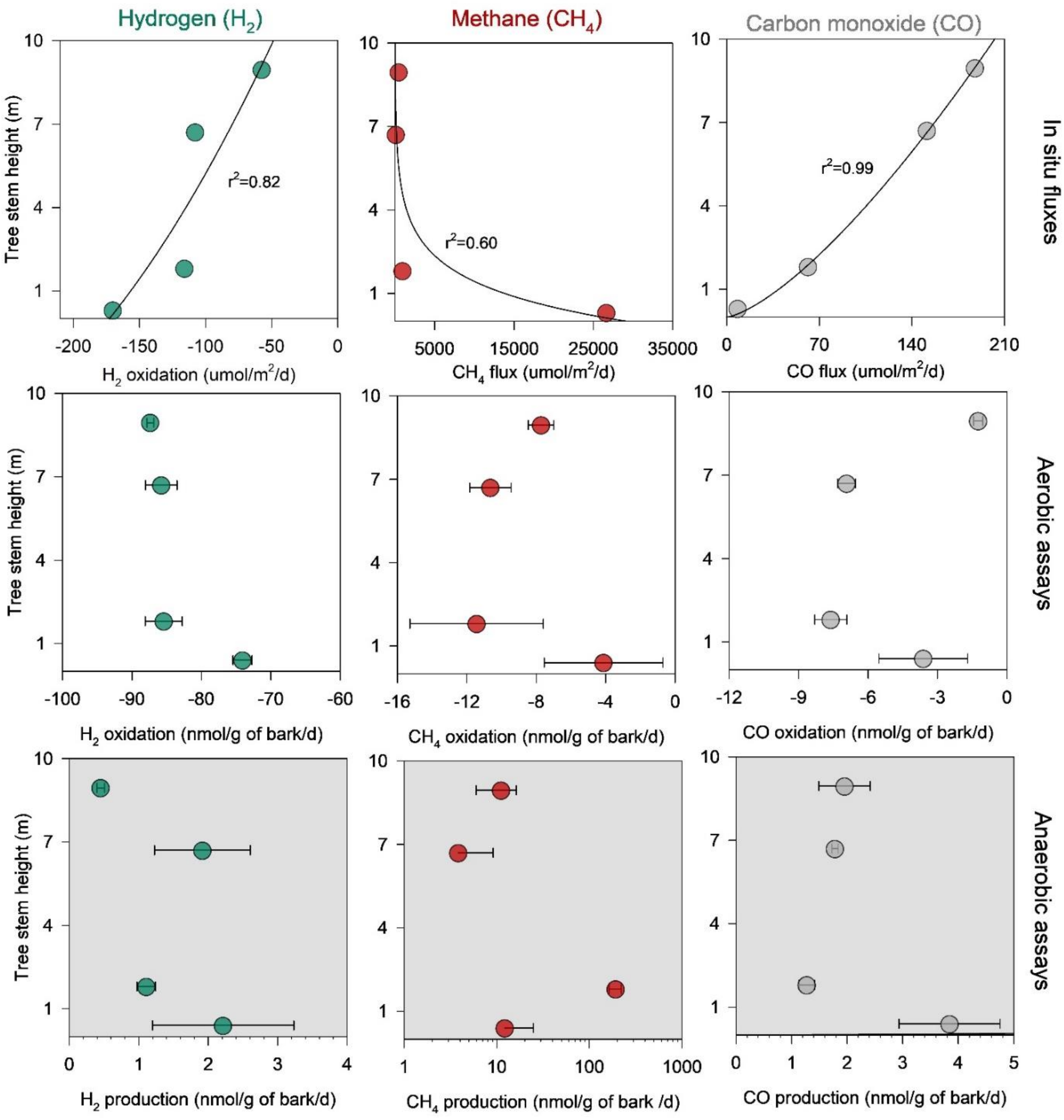

**Figure S8. *In situ* trace gas flux measurements**. Trace gas *M. quinquenervia* tree stem flux incubations showing the incremental changes of CO, $CH_4$, and $H_2$ during dry (left) vs wet (right) field conditions. Note: different log scale for $CH_4$ between campaigns.

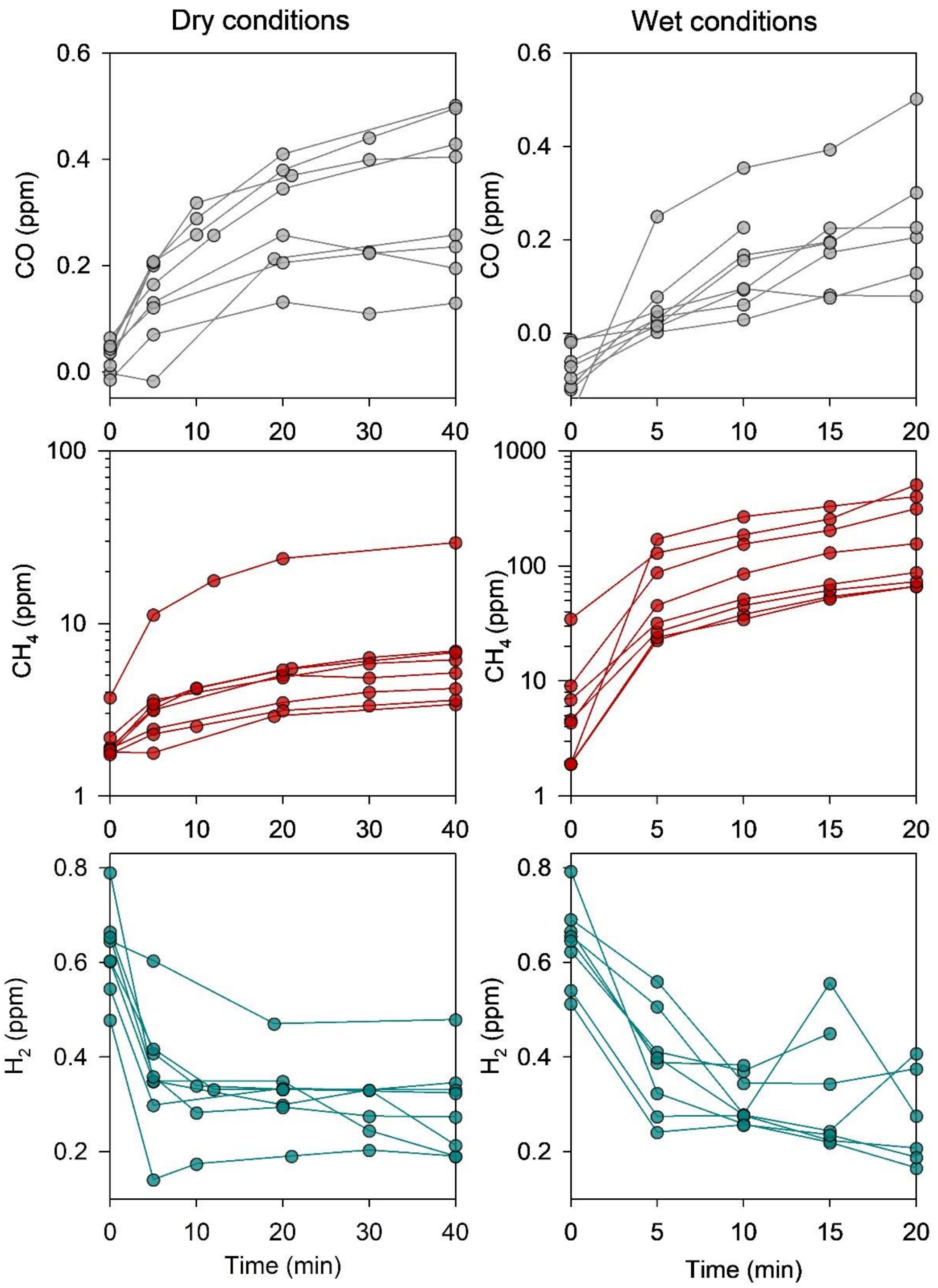

### Dataset

**Dataset S1 (xlsx)**. Sample metagenome metadata and 16S rRNA gene copy numbers.

**Dataset S2 (xlsx)**. Microbial community composition based on SingleM.

**Dataset S3 (xlsx)**. Relative abundance of metabolic marker genes in metagenome short reads.

**Dataset S4 (xlsx)**. Summary of taxonomy, quality statistics, coverage, derived metabolic gene protein sequences, and genetic capabilities of the metagenome-assembled genomes.

**Dataset S5 (xlsx)**. Summary of derived metabolic gene protein sequences in unbinned contigs.

**Dataset S6 (xlsx)**. Microbial community composition based on phyloFlash.

**Dataset S7 (xlsx)**. Summary of 47 publicly available tree species genomes for host sequence removal.
